# cGASylation by a bacterial E1-E2 fusion protein primes antiviral immune signaling

**DOI:** 10.1101/2022.03.31.486616

**Authors:** Hannah E. Ledvina, Qiaozhen Ye, Yajie Gu, Yun Quan, Rebecca K. Lau, Huilin Zhou, Kevin D. Corbett, Aaron T. Whiteley

**Affiliations:** Department of Biochemistry, University of Colorado Boulder, Boulder, Colorado, USA; Department of Cellular and Molecular Medicine, University of California, San Diego, La Jolla, California, USA; Biomedical Sciences Graduate Program, University of California, San Diego, La Jolla, California, USA; Department of Chemistry, University of California, San Diego, La Jolla, California, USA

## Abstract

In all organisms, innate immune pathways sense viral infection and rapidly activate potent immune responses while maintaining a high degree of specificity to prevent inappropriate activation (autoimmunity). In humans, the innate-immune receptor cGAS detects viral infection to produce the nucleotide second messenger cGAMP, which initiates STING-dependent antiviral signaling. Bacteria encode predecessors of the cGAS-STING pathway, termed cyclic oliogonucleotide-based antiphage signaling systems (CBASS), and bacterial cGAS detects bacteriophage infection to produce cGAMP. How bacterial cGAS activation is controlled, however, remains unknown. Here, we show that the CBASS-associated protein Cap2 primes bacterial cGAS for activation through a ubiquitin transferase-like mechanism. A cryoelectron microscopy structure of the Cap2–cGAS complex reveals Cap2 as an all-in-one ubiquitin transferase-like protein, with distinct domains resembling the eukaryotic E1 protein ATG7 and the E2 proteins ATG10 and ATG3. The structure captures a reactive-intermediate state with the cGAS C-terminus extending into the Cap2 E1 active site and conjugated to AMP. We find that Cap2 ligates the cGAS C-terminus to a target molecule in cells, a process we call cGASylation. cGASylation primes cGAS for a ∼50-fold increase in cGAMP production. We further demonstrate that Cap2 activity is balanced by a specific endopeptidase, Cap3, which deconjugates cGAS and antagonizes antiviral signaling. Our data demonstrate that bacteria control immune signaling using an ancient, minimized ubiquitin transferase-like system and provide insight into the evolution of E1 and E2 machinery across the kingdoms of life.

---

Antiviral innate immune pathways across life must rapidly sense and respond to a wide array of viral threats while limiting their activation in the absence of stimulus, which could otherwise lead to autoimmune disease or premature cell death. In eukaryotes, viral defense is in part mediated by the cGAS-STING (cyclic GMP–AMP Synthase-Stimulator of Interferon Genes) pathway^1^. Signaling is initiated when cGAS senses infection and generates the second messenger 2′,3′ cyclic GMP–AMP, which in turn binds to and activates STING to initiate a potent interferon response^2^. Recent work has demonstrated that the cGAS-STING pathway originated from bacterial cyclic oliogonucleotide-based antiphage signaling systems (CBASS), where they serve an analogous function in the bacterial antiviral immune response^3–6^.

CBASS antiviral pathways are widespread in bacteria and protect populations from phage infection by triggering programmed cell death^3,7,6^. CBASS operons encode a core cGAS/DncV-like nucleotidyltransferase (CD-NTase, here termed cGAS) that is activated upon phage infection^3^ and synthesizes one of a variety of cyclic di- and trinucleotide second messenger molecules^8^. Those molecules in turn activate a cell-killing effector protein^9,10^ to halt phage replication, a process termed abortive infection^3,4,11^. CBASS operons are classified based on operon architecture^6,7^. Type I CBASS systems encode only a cGAS enzyme and an effector protein, while the majority of CBASS systems (Types II, III, and IV) also encode additional proteins with proposed roles in phage sensing and cGAS activation. The mechanisms underlying CBASS-induced cell death have recently been the focus of numerous studies^4,10,12^, but the role of the additional proteins remains largely unknown.

We focused on Type II CBASS, which make up over 40% of total CBASS systems^7^, and selected a representative system from *Vibrio cholerae* that emerged in the genome of strains that caused the 7^th^ cholera pandemic (**Fig. 1a)**^13,14^. Upon phage infection, cGAS (also called DncV in this system) synthesizes the cyclic dinucleotide second messenger 3′,3′ cyclic GMP– AMP (cGAMP)^3,13^, which activates a cell-killing phospholipase effector (CapV)^9^. The *dncV*-*capV* operon, like all Type II CBASS, encodes two additional proteins termed CD-NTase-associated proteins 2 and 3 (Cap2 and Cap3)^7^, whose functions are unknown (**Fig. 1a**). When expressed in *Escherichia coli, V. cholerae* CBASS confers broad resistance to phage infection (**Fig. 1b, Extended data Fig. 1a**). Consistent with previous reports^3^, we found that cGAS, CapV, and Cap2 are all required for resistance, while Cap3 is dispensable (**Fig. 1b, Extended data Fig. 1a**).

**Figure 1:**
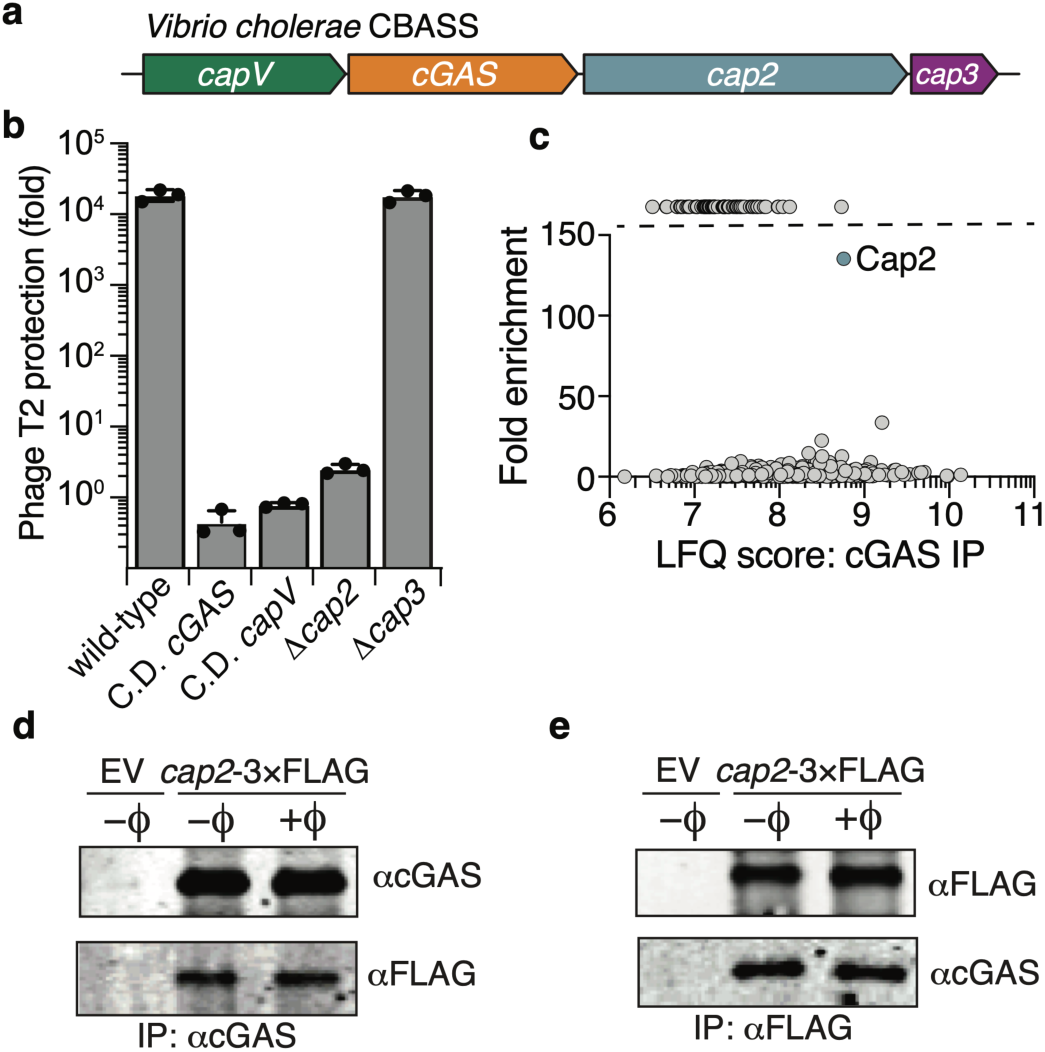
Cap2 is essential for CBASS function and directly interacts with cGAS. **(a)** Operon structure of CBASS from *V. cholerae*; see **Supplementary Table 7** for relevant accession numbers. **(b)** Efficiency of plating of phage T2 when infecting *E. coli* expressing *V. cholerae* CBASS with the indicated genotype. Data represent fold decrease in plaque forming units compared to bacteria expressing an empty vector. Catalytically dead (C.D.) cGAS: DID131AIA.; C.D. *capV*: C62S. **(c)** Mass spectrometry of immunoprecipitated cGAS. Data are label-free quantitation (LFQ) score and fold enrichment comparing immunoprecipitations from bacteria expressing wild-type CBASS to an empty vector control. Cap2, is represented by an orange circle and is labeled. **(d)** Western blot analysis of αcGAS immunoprecipitation from *E. coli* expressing empty vector (EV) or CBASS with the indicated genotype. ±ϕ indicates phage T2 at a multiplicity of infection (MOI) of 2. **(e)** Western blot analysis of αFLAG immunoprecipitation from *E. coli* expressing empty vector (EV) or CBASS with the indicated genotype. ±ϕ indicates phage T2 at an MOI of 2.

To understand how cGAS is regulated during infection, we immunoprecipitated cGAS from phage-infected bacteria (**Extended data Fig. 1b)**. Mass spectrometry revealed that Cap2 copurified with cGAS, suggesting these two proteins form a complex (**Fig. 1c, Supplementary Table 1**). Using a functional C-terminal 3×FLAG tag on Cap2 and reciprocal immunoblots, we confirmed a direct association between Cap2 and cGAS (**Fig. 1d-e, Extended data Fig 1c**). Unexpectedly, we found that the Cap2–cGAS interaction is independent of phage infection (**Fig. 1d-e**). Taken together with the essential role for Cap2 in protection against phage (**Fig. 1b**), these results suggest that Cap2 may be important for cGAS regulation prior to infection.

## Structure of the Cap2–cGAS complex

To understand the function of Cap2 and characterize the molecular basis for its cGAS interaction, we purified a stoichiometric Cap2–cGAS complex from a related CBASS found in *Enterobacter cloacae* (**Fig. 2a**) and determined a 2.7 Å-resolution structure by cryoelectron microscopy (cryoEM; **Fig. 2b, Extended Data Fig. 2, Supplementary Table 2**). The structure reveals a 2:2 complex with a homodimer of Cap2 bound to two cGAS monomers (**Fig. 2c**). Cap2 adopts a modular architecture with three domains: an N-terminal E2 domain, a central linker domain, and a C-terminal adenylation/E1 domain **(Fig. 2a)**. E1 and E2 domains catalyze post-translational modifications. The most frequent modification is transfer of ubiquitin or a related β-grasp fold protein (collectively termed ubiquitin-like proteins or Ubls) to target molecules^15^. In these pathways, E1 proteins first conjugate adenosine monophosphate (AMP) to the Ubl C-terminus (adenylation), then form a thioester bond between the Ubl C-terminus and the catalytic cysteine of the E1 protein. The Ubl is next shuttled to a cysteine residue on E2, and finally the Ubl is transferred to a target, often a lysine residue side chain or the N-terminus of a target protein, with the help of an E3 protein that typically serve as an adapter^16^.

**Figure 2:**
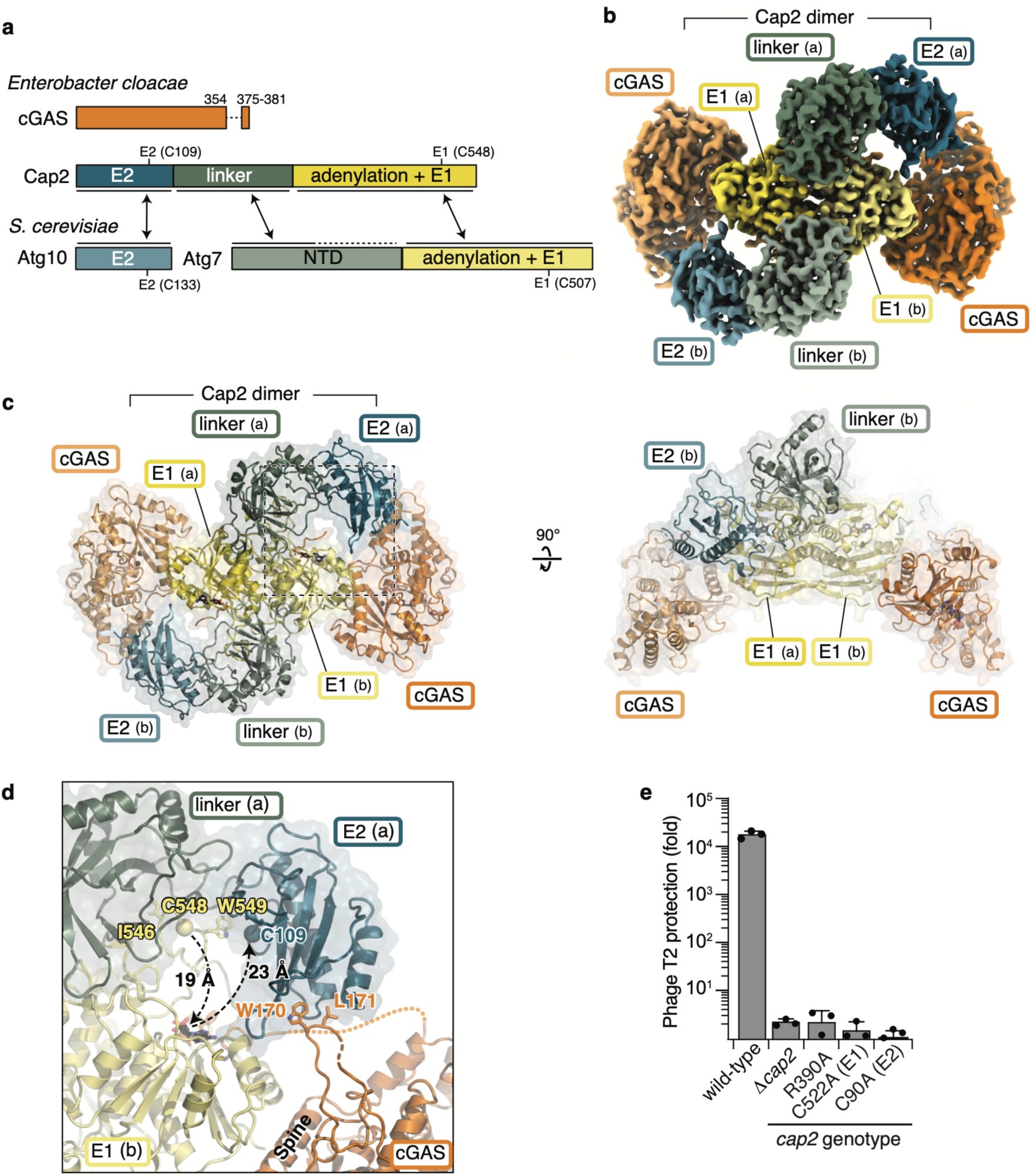
Cryoelectron microscopy structure of a Cap2–cGAS complex. **(a)** Domain schematic of cGAS and Cap2 from *Enterobacter cloacae* and Atg10 and Atg7 from *Saccharomyces cerevisiae*, with domains colored and labeled to represent similarity. See also **Extended Data Fig. 3. (b)** 2.74 Å resolution cryoEM density for the *E. cloacae* Cap2–cGAS(C019A/C548A) complex, with domains colored as in (a). See also **Supplementary Table 2** and **Extended Data Fig. 2. (c)** Two views of the 2:2 heterotetrameric Cap2–cGAS complex, with domains colored as in (a). **(d)** View of one set of active sites in the Cap2–cGAS complex, with the E1 and E2 active-site residues (C548 and C109, respectively; both mutated to alanine in this structure) shown as spheres. **(e)** Efficiency of plating of phage T2 when infecting *E. coli* expressing CBASS with the indicated genotype. Data plotted as in Fig. 1b.

In Cap2, the C-terminal adenylation/E1 domain forms a tight homodimer, similar to that observed in the bacterial E1 proteins MoeB and ThiF, which participate in sulfur metabolism rather than protein post-translational modification^16–22^. The central linker domain of each Cap2 protomer reaches over the E1 domain of its dimer mate, positioning each N-terminal E2 domain close to the active site of the dimer-related E1 domain (**Fig. 2c-d**). Each Cap2 protomer is bound to a monomer of cGAS via the underside of its E1 domain, with a second smaller interface between the nearby dimer-related E2 domain and an extended loop unique to cGAS enzymes in Type II CBASS systems (**Fig. 2c-d, Supplementary Table 3**^5,8,23,24^). Mutations in these interfaces reduce or eliminate Cap2 binding to cGAS (**Extended Data Fig. 3a-b**). The bipartite Cap2–cGAS interaction appears to rigidify Cap2 and position the E2 domain: a second structure from the same cryoEM dataset shows that when the Cap2 dimer is bound to only one cGAS monomer, the linker and E2 domains of the unbound Cap2 protomer become more flexible (**Extended Data Fig. 3c-d**).

The overall structure of Cap2 is more similar to the non-canonical E1 and E2 signaling machinery involved in autophagy than to the canonical systems that mediate ubiquitination (**Extended Data Fig. 4**). ATG7 is a eukaryotic non-canonical E1 enzyme that cooperates with the non-canonical E2 enzymes ATG3 or ATG10 to mediate transfer of the Ubls ATG8 (also called LC3 in humans) and ATG12^15,16^. ATG10 coordinates transfer of ATG12 to the target ATG5. ATG3, on the other hand mediates transfer of ATG8 to a phospholipid. ATG7, ATG3, and ATG10 have key structural differences compared to canonical E1 and E2 proteins. ATG7 forms a homodimer through its C-terminal E1 domain and possesses an N-terminal domain that recruits and positions E2 for catalysis^25–27^ (**Extended Data Fig. 4a-c**). Further, ATG7 possess a catalytic cysteine residue in the “crossover loop” that extends over the E1 adenylation active site, rather than within an α-helical insertion into this loop as in canonical E1 proteins^28,29^ (**Extended Data Fig. 4a-b, 5**). ATG3 and ATG10, lack conserved features of the canonical E2 proteins including an intact UBC fold and instead contain an extended hairpin loop on which the catalytic residue resides (**Extended Data Fig. 4d**). Consistent with Cap2 resembling ATG7, we see that Cap2 forms a homodimer through the E1 domain, possess a predicted catalytic cysteine on a crossover loop (C548), and encodes a linker domain that is homologous to ATG7 N-terminus that aides in coordinating the E2. In line with Cap2 resembling ATG3 and ATG10, we see that that the E2 domain possess homology to these proteins, including the location of the catalytic cysteine on an extended hairpin loop (C109, **Fig. 2d, Extended Data Fig. 4, 5**). The unambiguous similarity of Cap2 to ATG7, plus strong homology between the Cap2 E2 domain and ATG3 and ATG10, strongly suggest that these two systems share a common evolutionary origin and are distinct from eukaryotic canonical E1 and E2 proteins that catalyze ubiquitination or Ubl transfer (**Supplemental Discussion**). Our structure further suggests that Cap2 is an all-in-one transferase capable of protein ligation. Supporting this idea, disruption of the Cap2 adenylation active site or mutating the putative E1 or E2 catalytic cysteine residues to alanine eliminated the ability of *V. cholerae* CBASS to protect against phage infection (**Fig. 2e, Extended Data Fig. 6a**).

## Cap2 mediates cGASylation

Our structural and mutational data suggests that Cap2 possesses non-canonical E1 and E2 transferase activity essential for phage defense. However, CBASS operons do not encode a β-grasp fold protein like ubiquitin, therefore the substrate of Cap2 was unclear (here, “substrate” denotes the protein equivalent to ubiquitin in ubiquitination pathways, while “target” denotes the protein or other molecule to which the substrate is transferred). Based on the stable complex formed between Cap2 and cGAS *in vivo* (**Fig. 1d-e**), we hypothesized that cGAS is the Cap2 substrate. Despite the lack of sequence or structural homology between cGAS and ubiquitin, our structure of the Cap2–cGAS complex shows that the extreme C-terminus of cGAS (residues 375-381) is bound to the Cap2 E1 domain’s adenylation active site and conjugated to an AMP molecule (**Fig. 2d, 3a-b**). We confirmed this finding with a 2.1 Å-resolution x-ray crystal structure of the Cap2 E1 domain bound to a cGAS–AMP conjugate (**Extended Data Fig. 7, Supplementary Table 4**). The reactive-intermediate state captured in both structures closely matches prior structures of activated Ubls bound to their cognate E1 proteins^17,30,31^, consistent with cGAS serving as the substrate of Cap2. Further supporting our hypothesis, cGAS enzymes in Type II CBASS possess an extended, disordered C-terminal region with a highly conserved C-terminus that is similar to the conserved C-terminal diglycine motif of ubiquitin and related proteins (**Fig. 3c, Supplementary Table 3, Extended Data Fig. 8**).

**Figure 3:**
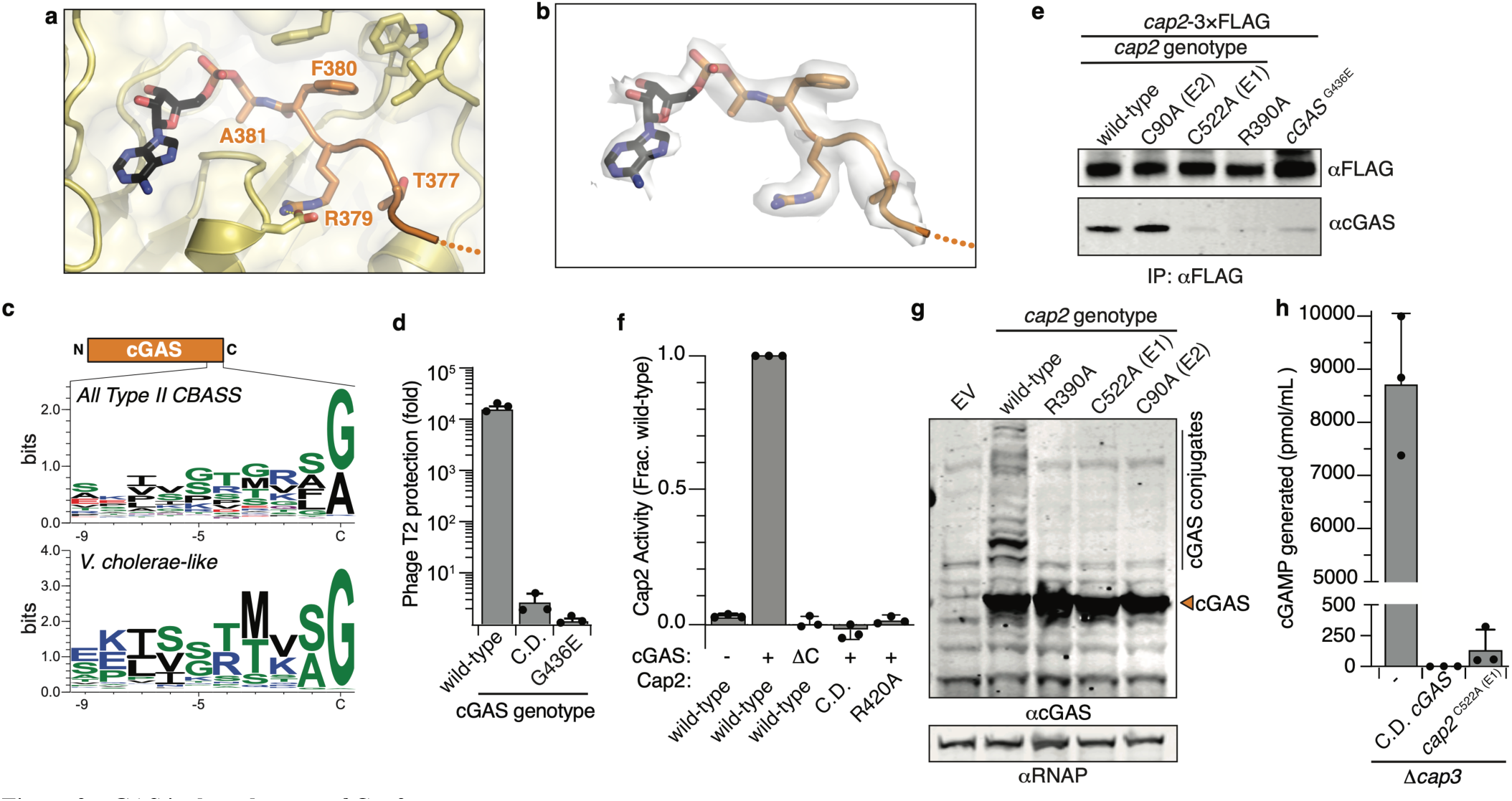
cGAS is the substrate of Cap2. **(a)** Cap2 adenylation active site in Cap2–cGAS cryoEM structure. cGAS C-terminus (orange) conjugated to AMP (black). **(b)** View equivalent to (a) showing unsharpened cryoEM density carved with a 2 Å radius around the cGAS–AMP conjugate. **(c)** Sequence logo for the C-terminal 10 residues of all 1556 cGAS enzymes from Type II CBASS or cGAS from only the *V. cholerae*-like group. Data is depicted as bits signified by the height of each residue. See also **Extended Data Fig. 8. (d)** Efficiency of plating by phage T2 when infecting *E. coli* expressing CBASS with the indicated genotype. Data plotted as in **Fig. 1b**. C.D. cGAS: DID131AIA. **(e)** Western blot analysis of anti-FLAG immunoprecipitation from *E. coli* expressing CBASS with the indicated genotype. **(f)** *E. cloacae* Cap2 activity assay, representing Cap2-mediated catalysis as a fraction of wild-type. The indicated genotypes of Cap2 and cGAS were expressed from a single plasmid and the formation of a cGAS conjugate was measured (in this assay, cGAS is conjugated to the flexible N-terminus of His6-Cap2; see **Extended Data Fig. 9** for details). (-): no cGAS; (+): wild-type cGAS; (ΔC): cGAS lacking its C-terminal 19 residues; C.D.: C548A/C109A catalytically dead Cap2. See **Extended Data Fig. 5** for Cap2 protein alignment. **(g)** Western blot analysis of cell lysates from *E. coli* expressing empty vector (EV) or CBASS with the indicated genotype. **(h)** cGAMP generated by cGAS immunoprecipitated from *E. coli* expressing CBASS with the indicated genotype. C.D. cGAS: DID131AIA catalytic- dead mutant.

If cGAS is the substrate of Cap2, mutating or deleting the disordered C-terminal region of cGAS or mutating the Cap2 E1 active site should destabilize the Cap2–cGAS complex and disrupt CBASS signaling. Accordingly, mutation of the C-terminal glycine residue of *V. cholerae* cGAS to a bulky, negatively-charged glutamate residue (G436E) resulted in loss of phage protection (**Fig. 3d**). We further found that the Cap2– cGAS interaction in bacterial cells is compromised upon mutation of the Cap2 adenylation active site, the E1 catalytic cysteine, or the C-terminus of cGAS (**Fig. 3e, Extended Data Fig. 6b-c**). In parallel, we established an in-cell Cap2 activity assay and found that, indeed, Cap2-mediated cGAS conjugation depends on the Cap2 adenylation active site, E1 and E2 catalytic cysteines, and on the cGAS C-terminus (**Fig. 3f, Extended Data Fig. 9**). Finally, we hypothesized that if the cGAS C-terminus is being conjugated to target molecules, then we should be able to identify changes in the apparent molecular weight of cGAS in denaturing SDS-PAGE analysis. Consistent with this hypothesis, we observed a series of novel, high-molecular-weight bands in an anti-cGAS immunoblot from CBASS-expressing bacteria, which disappear when the catalytic functions of Cap2 are disrupted (**Fig. 3g**). In total these data demonstrate that Cap2 is a bona fide cGAS transferase, catalyzing a process we term cGASylation.

## cGASylation increases the enzymatic activity of cGAS

To directly test the functional impact of cGASylation, we immunoprecipitated cGAS from bacterial cells expressing wild-type and *cap2* mutant CBASS and measured synthesis of the second messenger cGAMP. We found that cGAS purified from cells with wild-type Cap2 was ∼50-fold more catalytically active than cGAS purified from cells with a Cap2 E1 active-site cysteine mutant (**Fig. 3h**). These data demonstrate that cGAMP synthesis by cGAS is significantly enhanced by cGASylation. The relevant target protein or molecule modified by cGASylation remains unknown and will be the subject of future studies. However, these data demonstrate that Cap2-mediated cGASylation dramatically increases the enzymatic potential of cGAS and provide a mechanism for why Cap2 is required for phage resistance.

## Cap3 antagonizes Cap2 by cleaving cGAS-target conjugates

All CBASS systems that encode Cap2 also encode Cap3, which is homologous to eukaryotic JAB/JAMM-family deubiquitinases^6,7^ (**Fig. 1a, Extended Data Fig. 11a**,**d)**. While Cap3 is dispensable for phage resistance (**Fig. 1b**), we hypothesized that this protein may balance CBASS activation by proteolytically cleaving cGAS–target conjugates. We tested this hypothesis by overexpressing Cap3 during infection and found that expression of wild-type Cap3, but not a catalytically-dead mutant, strongly counteracted CBASS-mediated defense (**Fig. 4a, Extended Data Fig. 10**). To directly measure Cap3 activity, we incubated purified *V. cholerae* Cap3 with a model substrate comprising cGAS fused at its C-terminus to GFP. Cap3 specifically removed cGAS from GFP *in vitro*, and this activity was disrupted by mutation of the Cap3 putative active site or chelation of the critical active-site Zn^2+^ ion by EDTA (**Fig. 4b, Extended Data Fig. 10a**). Mass spectrometry revealed that Cap3 cleaves this model substrate specifically after the C-terminal residue of cGAS (**Fig. 4c, Extended Data Fig. 11b, Supplementary Table 5**). Cap3 was unable to cleave substrates with mutations in the extreme C-terminal region of cGAS, demonstrating a high degree of specificity (**Extended Data Fig. 11c**). We observed similar results with *E. cloacae* Cap3, including specific cleavage at the C-terminus of *E. cloacae* cGAS that was disrupted by mutations to the cGAS C-terminal region (**Extended Data Fig. 11d-g**).

**Figure 4:**
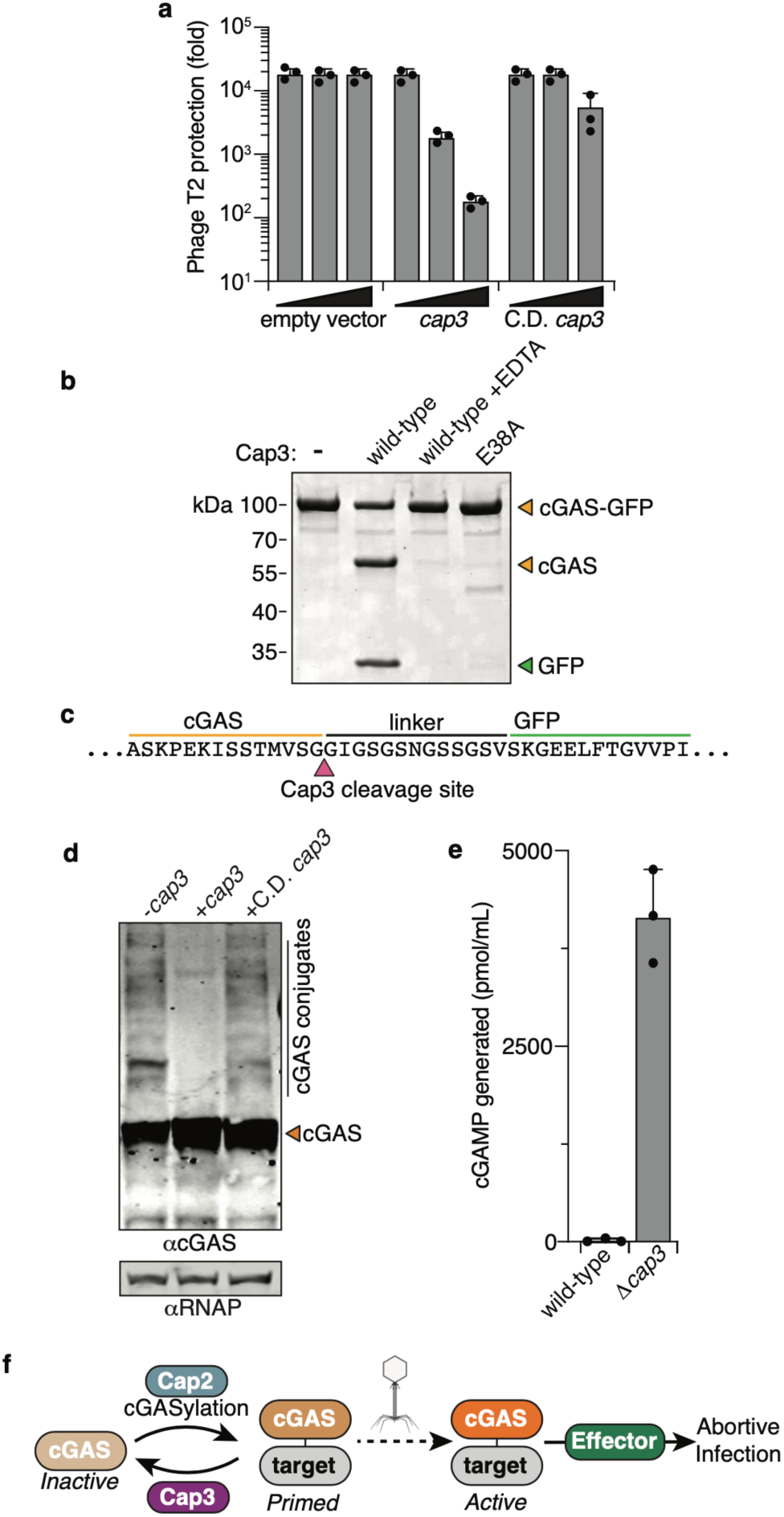
Cap3 antagonizes CBASS signaling by cleaving products of cGASylation. **(a)** Efficiency of plating of phage T2 when infecting *E. coli* expressing CBASS Δ*cap3* in the absence or presence of overexpressed *cap3* with the indicated genotype. Data plotted as in **Fig. 1b**. C.D. *cap3*: HTH101ATA. See **Extended Data Fig. 10** for protein alignment. **(b)** Coomassie stained SDS-PAGE of an *V. cholerae* model substrate (cGAS-GFP fusion protein) incubated with *V. cholerae* Cap3 with the indicated reaction condition/genotype. **(c)** Summary of tryptic digest mass spectrometry analysis of the Cap3-treated cGAS bands as in (b), showing the putative *V. cholerae* Cap3 cleavage site. See **Supplementary Table 5** for full data. **(d)** Western blot analysis of cell lysates from *E. coli* expressing CBASS Δ*cap3* plus a second vector expressing *cap3* with the indicated genotype. C.D. *cap3*: HTH101ATA. **(e)** cGAMP generated by cGAS immunoprecipitated from *E. coli* expressing CBASS with the indicated genotype. **(f)** Model depicting the role of Cap2 and Cap3 in CBASS regulation. Briefly, Cap2 conjugates cGAS to an unknown target via E1-E2 ubiquitin transferase-like mechanism, a process here termed cGASylation. cGASylation primes cGAS for activation by phage infection. Upon infection, cGAS becomes enzymatically active and generates a cyclic oligonucleotide second messenger which then activates an effector protein. Effector protein activity leads to cell death, inhibiting phage by abortive infection. cGASylation is antagonized by Cap3 protease activity which removes cGAS from target proteins, thereby limiting cGAS priming.

Finally, we tested the ability of Cap3 to antagonize cGASylation in cells. Overexpression of wild-type Cap3, but not a catalytic-dead mutant, eliminated the formation of Cap2-dependent high molecular weight cGAS bands (**Fig. 4d**). Further, Cap3 reduced cGAS-mediated cGAMP synthesis by over 200-fold compared to cells without Cap3 (**Fig. 4e**). These data strongly support a model in which cGAS activation is potentiated by cGASylation, and that Cap3 antagonizes this activity by cleaving cGAS–target conjugates (**Fig. 4f**).

## The multifaceted regulation of CBASS

A recent renaissance in the study of phage resistance has revealed many novel antiviral innate immune systems, sometimes encoding protein domains and components that are structurally and functionally conserved across domains of life^3,5,32–34^. While significant progress has been made in identifying these systems^33,35^, their molecular mechanisms remain unclear. In this study, we show that the CBASS protein Cap2 is structurally homologous to ubiquitin transferases and most closely resembles non-canonical E1 and E2 transferases associated with autophagy in eukaryotes. Cap2 conjugates bacterial cGAS to target molecules in a process we term cGASylation. The covalent cGAS adduct is primed for activation and essential for phage defense, but can be reversed by Cap3, a sequence-specific protease (**Fig. 4f)**. Cap2, like other CBASS genes, is evolutionarily related to key components of the eukaryotic immune system^3,5,36^. Ubiquitin transferases and subsequent signaling play a central role in human innate immune pathways, such as TLR3/4 and RIG-I, by mediating protein-protein interactions required for signaling^37–39^. Taken together with our findings, E1 and E2-domain mediated protein conjugation represent a conserved mechanism of immune regulation.

While bacterial ubiquitin transferases have been identified^40^, our structural and biochemical findings reveal Cap2 as the first known all-in-one ATP-dependent ubiquitin transferase-like protein that combines adenylation, E1, and E2 active sites into a single polypeptide. Based on the lack of an E3 protein in Type II CBASS operons and bacteria, we hypothesize that target recognition is mediated directly by Cap2. The striking similarity of Cap2 to non-canonical E1 and E2 transferases from eukaryotes (ATG7 and ATG3/ATG10) suggests that these systems share a common evolutionary origin. Thus, while common ancestors of canonical ubiquitin signaling can be found throughout eukaryotes and in some archaea^41,42^, E1 and E2 transferases may have evolved first in bacteria, in line with prior bioinformatic observations^19,21^. Our work also reveals bacterial cGAS enzymes are the first known substrates of ubiquitin transferase-like systems that do not share the β-grasp fold of ubiquitin and related proteins^43^. The all-in-one nature of Cap2, and its unique mode of substrate recognition, may enable engineering of this system to mediate customizable post-translational modifications, including scar-free protein fusions generated either *in vitro* or *in vivo*. Similarly, a complete understanding of Cap3 specificity will enable future applications of these proteases for site-specific cleavage of peptide and isopeptide bonds.

What is the role of cGASylation in CBASS signaling? We hypothesize that Cap2 primes cGAS for activation and cGAMP production, and that cGAS is the “phage sensor” whose direct stimulus remains unknown. This is consistent with the emerging general model for CBASS signaling, in which cGAS activity is held in check by inhibitory molecules depleted during phage infection^3,8,10,42^. cGASylation is not altered by phage infection but instead may prime the system to increase sensitivity, enabling cGAS to quickly respond to phage infection. Alternatively, cGASylation may license cGAS to counter inappropriate activation. The abundance of primed cGAS is further tuned by the activity of Cap3, which antagonizes cGASylation, striking a balance between activation sensitivity and specificity that is reminiscent of human innate-immune pathways that must be balanced to avoid autoimmune reactions^37,46,47^. Taken together, our results demonstrate that Type II CBASS systems use unique regulatory mechanisms reminiscent of ubiquitin-mediated immune regulation in humans to finely control the activation of cGAS enzymes and mediate broad antiphage activity.

## Supplemental discussion

The evolutionary origin of eukaryotic ubiquitin signaling pathways has long been a topic of intense interest. All E1 proteins likely evolved from bacterial enzymes related to the homodimeric E1s MoeB and ThiF, which adenylate the C-terminus of the ubiquitin-like proteins (Ubls) MoaD and ThiS, respectively, in sulfur-transfer metabolic pathways (reviewed in ^16,21,67,68^). Also in bacteria, several families of operons encoding different combinations of Ubl, E1, E2, and JAB peptidase have been identified but not functionally characterized, leaving open the question of whether these systems mediate protein transfer^21,43^. Five distinct bacterial operon types have been described (termed families 6a-6e), of which Cap2 and Cap3-containing CBASS systems are one (family 6b)^43^.

Recent data have shown that many archaea possess E1, E2, and RING-family E3 ligases that together conjugate a Ubl to target proteins, providing strong evidence that eukaryotic ubiquitin signaling evolved from these archaeal systems^41,42,69,70^. The evolutionary origin of the non-canonical eukaryotic E1 and E2 proteins ATG7, ATG3, and ATG10, which conjugate the Ubls ATG12 and ATG8 to a target protein (ATG5) and a phospholipid (phosphatidylethanolamine), respectively^71^, are less well defined.

Several lines of evidence support an evolutionary relationship between Cap2 and the autophagy-related non-canonical E1 (ATG7) and E2 (ATG3 and ATG10) proteins. First, unlike most eukaryotic E1s that contain a single active adenylation site, the C-terminal E1 domain of Cap2 forms a homodimer with two active adenylation sites, like MoeB^17^, ThiF^22^, and ATG7^25–27^ (**Extended Data Fig. 4a-b**). The ATG7 E1 domain is also more closely related to bacterial E1s in the 6a and 6b/Cap2 families than it is to other eukaryotic E1 proteins^21,43,70^. Second, while the catalytic cysteine residues of most eukaryotic E1 proteins are located on an α-helical domain inserted in the crossover loop of their adenylation domain^16^, the catalytic cysteines of both Cap2 and ATG7 are located on these proteins’ crossover loop, with no α-helical domain insertion (**Extended Data Fig. 4b**). Thus, Cap2 and ATG7 share a common structural mechanism of substrate adenylation and transfer to the E1 catalytic cysteine.

In most eukaryotic E1 proteins, a ubiquitin fold domain (UFD) positioned C-terminal to the adenylation domain recruits and positions E2 proteins for substrate transfer^16^. In ATG7, however, the protein’s structurally distinctive N-terminal domain recruits and positions two different non-canonical E2 proteins, ATG3 and ATG10, for catalysis^28,29^. The central linker domain of Cap2 shares a common fold with the ATG7 N-terminal domain (**Extended Data Fig. 4c**) and is positioned similarly in the dimeric assembly, with each linker domain draping over its dimeric partner’s adenylation active site (**Extended Data Fig. 4b**). While the ATG7 N-terminal domain recruits separate E2 proteins, the Cap2 linker domain positions its own N-terminal E2 domain close to the adenylation site and E1 catalytic cysteine. Thus, the role of this domain in positioning E2 for catalysis is shared between Cap2 and ATG7.

In addition to similarities between Cap2 and ATG7, the Cap2 N-terminal E2 domain is structurally related to the non-canonical E2 proteins ATG3 and ATG10 that play a role in autophagy. Canonical E2 proteins contain a UBC fold, which comprises a four-stranded β-sheet surrounded by four α-helices, with the catalytic cysteine located on a loop between β4 and α2^16^. Like ATG3 and ATG10, the Cap2 E2 domain lacks α-helices 3 and 4 (**Extended Data Fig. 4d**), and also lacks other conserved features of canonical E2 proteins, including an HPN motif ∼8 amino acids N-terminal to the catalytic cysteine, a conserved tryptophan residue C-terminal to the catalytic cysteine, and a proline-rich motif between β-strands 3 and 4^72^. Further, the catalytic cysteine of the Cap2 E2 domain (C109) is positioned on a long hairpin loop that is structurally analogous to the extended β5-β6 hairpin found in both ATG3 and ATG10 (**Extended Data Fig. 4d**).

Cap2 possesses strong structural similarity to the autophagy E1 ATG7 and to its cognate E2s ATG3 and ATG10. Thus, despite important differences between Cap2 and the autophagy E1/E2 proteins, including distinct substrates (cGAS versus Ubl proteins) and protein architecture (a single polypeptide versus separate E1 and E2 proteins), we conclude that these pathways share a common bacterial ancestor distinct from the archaeal ancestors of other eukaryotic Ubl transfer pathways.

## Acknowledgements

The authors thank R. Parker and A. Desai for critical reading of the manuscript; M. Herzik for advice on cryoEM data collection and processing; C. Ebmeirer and the Mass Spectrometry Facility at the University of Colorado Boulder (RRID:SCR_018992) for assistance with sample preparation, experimental details, and data analysis (NIH S10-OD025267); the CU Boulder Department of Biochemistry Shared Instruments Pool core facility (RRID:SCR_018986) and its staff; J. Kralj at CU Boulder for use of his laboratory’s plate reader; and members of the Corbett and Whiteley labs for advice and helpful discussion. The authors acknowledge the facilities of the cryoEM facility at UC San Diego, and technical assistance of R. Ashley on cryoEM sample preparation and data collection. This work was funded by NIH R21AI148814 (K.D.C.), a Mallinckrodt Foundation Grant (A.T.W.), and NIH R01 GM116897 and S10 OD023498 (H.Z.). H.E.L. is supported as a fellow of the Jane Coffin Childs Memorial Fund for Medical Research and R.K.L. is supported by NIH F31GM137600. X-ray diffraction data were collected at the Northeastern Collaborative Access Team beamlines, which are funded by the National Institute of General Medical Sciences from the National Institutes of Health (P30 GM124165). The Eiger 16M detector on 24-ID-E is funded by a NIH-ORIP HEI grant (S10OD021527). This research used resources of the Advanced Photon Source, a U.S. Department of Energy (DOE) Office of Science User Facility operated for the DOE Office of Science by Argonne National Laboratory under Contract No. DE-AC02-06CH11357.

## Author contributions

Experiments were designed and conceived by H.E.L., Q.Y., A.T.W., and K.D.C. Phage assays, *in vivo* bacterial assays, and cGAMP generation assays were performed by H.E.L. Biochemical experiments were performed by Q.Y. Mass spectrometry experiments were performed by Q.Y., Y.Q., and H.E.L. Structural experiments and analysis were performed by Q.Y., Y.G., and K.D.C. Protein purification was performed by Q.Y. and R.K.L. Bioinformatic analysis of cGAS proteins was performed by K.D.C. Figures were prepared by H.E.L., A.T.W. and K.D.C. The manuscript was written by H.E.L., A.T.W., and K.D.C. All authors contributed to editing the manuscript and support the conclusions.

## Materials and Methods

### Bacterial strains and growth conditions

*E. coli* strains used in this study are listed in **Supplementary Table 6**. *E. coli* were cultured in LB medium (1% tryptone, 0.5% yeast extract, 0.5% NaCl) at 37 °C unless otherwise noted. For phage experiments and other noted assays, bacteria were grown in “MMCG” minimal medium containing M9 salts, magnesium, calcium, and glucose (47.8 mM Na_2_HPO4, 22 mM KH_2_PO_4_, 18.7 mM NH_4_Cl, 8.6 mM NaCl, 22.2 mM glucose, 2 mM MgSO_4_, 100 μM CaCl_2_, 3 μM Thiamine). Where applicable, media were supplemented with carbenicillin (100 μg/mL) or chloramphenicol (20 μg/mL), to ensure plasmid maintenance. When a strain with two plasmids was cultivated in MMCG medium, bacteria were cultured with 20 μg/mL carbenicillin and 4 μg/mL chloramphenicol. All strains were stored in LB plus 30% glycerol at −70 °C.

### Plasmid Construction

Plasmids used in the study are listed in **Supplementary Table 6**. All experiments were performed with either the CBASS system from *V. cholerae* C6706 (NCBI RefSeq NZ_CP064350.1) or *E. cloacae* (NCBI RefSeq NZ_KI973084.1; see protein accession numbers in **Supplementary Table 7**). For phage infections, the entire operon plus surrounding sequences were cloned into the XhoI and NotI sites of the vector pLOCO2 (Kibby et al., manuscript in preparation). For the *in vivo* Cap3 work, genes were cloned into the BamHI and NotI sites of the vector pTACxc. For immunoprecipitation assays, a C-terminal 3×FLAG tag was added to *V. cholerae cap2* and an N-terminal VSV-G tag was added to *V. cholerae* cGAS. For biochemical analysis, individual proteins were cloned into vector 2-BT (Addgene #29666; N-terminal His_6_-TEV cleavage site fusion), H6-msfGFP (Addgene #29725; N-terminal His_6_-TEV cleavage site fusion and C-terminal msfGFP fusion), or 2-AT (Addgene #29665; untagged).

For *E. cloacae* Cap3, sequence alignments revealed that the first 16 amino acids of the annotated gene are unlikely to be translated *in vivo*; a truncated construct comprising residues 17-180 of the annotated gene expressed at higher levels and was more soluble upon purification (for mutations, residue numbering follows the annotated gene). For *E. cloacae* Cap2-cGAS complex used for cryoEM, the two genes were PCR-amplified from vector 2-AT and combined to generate a polycistronic transcript, then cloned into vector 2-BT resulting in an N-terminal His_6_-tag on cGAS and no tag on Cap2, and both catalytic cysteine residues in Cap2 (C109 and C548) were mutated to alanine. For *E. cloacae* Cap2-cGAS complex used for N-terminal cGASylation, the two genes were cloned as above into vector 2-BT to generate a polycistronic transcript with an N-terminal His_6_-tag on Cap2 and no tag on cGAS. For *E. cloacae* Cap2-cGAS complex with HA-tagged cGAS, the two genes were cloned as above into vector 2-AT to generate a polycistronic transcript with an N-terminal HA tag (MYPYDVPDYAGSG) fused to residue 2 of cGAS.

DNA sequences were cloned into destination vectors using 18-25 bp overhangs and Gibson Assembly. Point-mutations and epitope tags were cloned by mutagenic PCR and isothermal assembly. Clones were transformed either into a modified strain of OmniMax *E. coli* (Invitrogen) by electroporation, or into NovaBlue *E. coli* (Novagen) by heat-shock and plated on LB with the appropriate selection. Positive clones were verified by Sanger Sequencing (Genewiz). Prior to use in downstream phage or immunoprecipitation experiments, sequence verified plasmids were transformed into MG1655 via heat shock and plated on LB with the appropriate selection.

### Phage Amplification and Storage

Phages used in the study are listed in **Supplementary Table 8**. Phage lysates were generated from *E. coli* MG1655 using a modified double agar overlay plate amplification (T2) or liquid amplification (T4, T5, T6). For plate amplification, stationary phase MG1655 was infected with 10,000 plaque forming units (PFU) of phage in LB + 0.35% agar, 10 mM MgCl_2_, 10 mM CaCl_2_, and 100 μM MnCl_2_. Plates were incubated overnight at 37 °C and the following day phages were harvested by adding 5 mL of SM buffer (100 mM NaCl, 8 mM MgSO_4_, 50 mM Tris-HCl pH 7.5, 0.01% gelatin) directly to the plate, incubating for 1 hour at room temperature, then collecting and filtering the resulting liquid through a 0.2 μm Nanosep filter. For liquid amplification, early logarithmic phase MG1655 was infected at an MOI of 0.1 in 25 mL LB broth plus 10 mM MgCl_2_, 10 mM CaCl_2_, and 100 μM MnCl_2_ at 37 °C with 220 rpm shaking for 2-6 hours until the culture became clear. Supernatants were then collected via centrifugation and filtration with a 0.2 μm Nanosep filter. Lysate titers were determined by spotting a serial dilution of the phage onto 0.35% LB agar plus 10 mM MgCl_2_, 10 mM CaCl_2_, and 100 μM MnCl_2_ containing stationary phase MG1655. Plates were incubated overnight at 37 °C and the resulting PFU/mL was calculated. Phage stocks were stored at 4 °C in either SM buffer or LB broth.

### Phage infection assays

Phage protection assays were performed by mixing 400 μL mid-logarithmic phase MG1655 containing the indicated vector(s) into 3.5 mL 0.35% MMCG agar and spotting a serial dilution of phage onto the agar. For the Cap3 overexpression experiments 0, 50, or 500 μM IPTG was added to both the bacterial culture and the top agar. After phage spots dried, plates were incubated at 37 °C overnight. The resulting PFU/mL was calculated, and data is presented as fold protection compared to a control strain expressing GFP. Data is shown as the mean +/-standard deviation of three biological replicates.

### Immunoprecipitation assays

MG1655 *E. coli* expressing the indicated vector(s) were grown to mid-logarithmic phase in MMCG. Where listed, cells were infected with the indicated phage for 30 minutes at an MOI of 2. Cultures were then centrifuged, and the resulting pellet was resuspended in lysis buffer (400 mM NaCl, 20 mM Tris-HCL pH 7.5, 2% glycerol, 1% triton, and 1 mM 2-mercaptoethanol). Cells were disrupted by sonication followed by centrifugation at 4 °C to remove cellular debris. Soluble lysates were then mixed with the affinity tag purification resin as described below overnight at 4 °C with end-over-end rotation. The following day, samples were washed five times in 1-5 mL lysis buffer and beads were processed for downstream application. For cGAS immunopreciptiations, lysates were incubated with protein A magnetic beads (Pierce) containing 10 μg/mL αcGAS antibody. Cap2-3×FLAG was immunoprecipitated using magnetic beads covalently linked to the αFLAG M2 antibody (Sigma). Finally, for the ELISA assays, VSV-G-cGAS was enriched through incubation with agarose beads conjugated to an αVSV-G antibody (Sigma).

### Western blots

αcGAS serum was generated by injecting rabbits with purified untagged cGAS, and polyclonal αcGAS antibodies were purified by antigen affinity (GenScript). Serum was used at 1:30,000 for cGAS detection. αFLAG antibody (Sigma) was used at 1:10,000 to detect Cap2-3×FLAG, αVSV-G (Rockland) was used at 1:7500 to detect VSV-G-cGAS, and αHA (clone 3F10, Sigma-Aldrich) was used at 1:30,000 to detect HA-tagged proteins.

For whole cell lysate analysis, 5 mL of MG1655 carrying the indicated plasmid were grown to mid-logarithmic phase. Cell densities were then normalized and 5×10^7^CFU were collected, centrifuged and resuspended in 50 μL of 1× LDS buffer (106 mM Tris-HCl pH7.4, 141 mM Tris Base, 2% w/v Lithium dodecyl sulfate, 10% v/v Glycerol, 0.51 mM EDTA, 0.05% Orange G). Samples were then incubated at 95 °C followed by a 5-minute centrifugation at 20,000×g to remove debris. For immunoprecipitation samples, affinity purification beads were resuspended in 40 μL lysis buffer plus 40 μL 2x LDS buffer. Samples were then incubated at 95 °C followed by a 5-minute centrifugation at 20,000×g.

Samples in LDS were loaded at equal volumes to resolve using SDS-PAGE, then transferred to PVDF membranes charged in methanol. Membranes were blocked in Licor Intercept Buffer for 30 minutes at 24 °C, followed by incubation with primary antibodies diluted in Intercept buffer overnight at 4 °C. Blots were then incubated with the appropriate combination of Licor infrared (800CW/680RD) αRabbit/Mouse secondary antibodies at 1:30,000 dilution in TBS-T (0.1% Triton-X) for 45 minutes at 24 °C and visualized using the Licor Odyssey CLx. For αHA immunoblots, HRP-linked Goat αRat antibody (Pierce 31470) was used at 1:30,000 and detected with an HRP Substrate kit (Bio-Rad) and Bio-Rad ChemiDoc imager. Representative images were assembled using Adobe Illustrator CC 2022.

### Mass Spectrometry analysis

Following IP enrichment as described above, samples were subjected to on bead trypsin digest followed by analysis on a Thermo Obitrap Q-Exactive HF-X using nanoLC-MS MS. Peptides were mapped to the proteome of *E. coli* MG1655 (https://www.uniprot.org/proteomes/UP000030788) plus the proteins composing the CBASS operon from *V. cholerae* (CapV, cGAS, Cap2, Cap3). The data point for cGAS is not shown in the figure for simplicity, however all data can be found in **Supplementary Table 1**.

### cGAS Enzyme Assay

6.25 × 10^9^ CFU of MG1655 expressing the indicated plasmids were processed for IP enrichments as described above using 15 μL bead volume. α-VSVG agarose beads were further washed three times in 1 mL reaction buffer (50 mM Tris-HCl pH 7.5, 50 mM KCl, 5 mM MgCl_2_). Samples were then resuspended in 50 μL reaction buffer and incubated with 500 μM ATP and 500 μM GTP overnight at 37 °C followed by 10 minutes at 95 °C to inactivate cGAS. cGAMP levels were then quantified with the Arbor Assay 3’,3’-cGAMP ELISA per manufacturer’s specifications. When necessary, samples were diluted 1:5 and 1:25 in the provided assay buffer to ensure measurements were in the dynamic range. cGAMP levels were calculated using the provided standard measured in triplicate. Data shown are a representative graph from one of three independent experiments depicting the mean +/- the standard deviation of three technical replicates.

### Protein Alignments

Cap2 and Cap3 protein alignments were generated with the MUSCLE algorithm^48^ within Geneious software, then adjusted by hand based on structure superpositions performed using the PDBeFold server (https://www.ebi.ac.uk/pdbe/).

### Protein Expression and Purification

Protein expression vectors used in the study are listed in **Supplementary Table 6**. For protein purification, expression vectors were transformed into *E. coli* Rosetta2 pLysS (EMD Millipore) or LOBSTR (Kerafast), grown at 37 °C in 2×YT media to an OD_600_ of 0.6, then protein expression was induced by the addition of 0.25 mM IPTG (isopropylthio-β-galactoside). Cultures were shifted to 20 °C for 16 hours, then cells were harvested by centrifugation. Cells were resuspended in binding buffer (25 mM Tris-HCl pH 8.5, 5 mM Imidazole, 300 mM NaCl, 5 mM MgCl_2_, 10% glycerol, and 5 mM 2-mercaptoethanol), lysed by sonication, and centrifuged (20,000 ×g for 30 minutes) to remove cell debris. Clarified lysate was passed over a Ni^2+^ affinity column (Ni-NTA Superflow, Qiagen) and eluted in a buffer with 250 mM imidazole. For cleavage of His_6_-tags, proteins were buffer-exchanged to binding buffer, then incubated 48 hours at 4 °C with His_6_-tagged TEV protease^49^. Cleavage reactions were passed through a Ni^2+^ affinity column again to remove uncleaved protein, His_6_-tags, and TEV protease. Flow-through fractions were passed over a size-exclusion chromatography column (Superdex 200; Cytiva) in gel filtration buffer (25 mM Tris-HCl pH 8.5, 300 mM NaCl, 5 mM MgCl_2_, 10% glycerol, 1 mM DTT). Gel filtration buffer without glycerol was used for samples for cryoelectron microscopy. Purified proteins were concentrated and stored at -80 °C for analysis or 4 °C for crystallization.

### Cryoelectron Microscopy

For grid preparation, freshly-purified *E. cloacae* Cap2-cGAS complex was collected from size-exclusion chromatography and diluted to 8 μM. Immediately prior to use, Quantifoil Cu 1.2/1.3 300 grids were glow-discharged for 10 sec in a pre-set program using a Solarus II plasma cleaner (Gatan). Sample was applied to a grid as a 3.5 μL drop in the environmental chamber of a Vitrobot Mark IV (Thermo Fisher Scientific) held at 4 °C and 100% humidity. After a 1-minute incubation, the grid was blotted with filter paper for 5 seconds prior to plunging into liquid ethane cooled by liquid nitrogen. Grids were mounted into standard AutoGrids (Thermo Fisher Scientific) for imaging.

All samples were imaged using a Titan Krios G3 transmission electron microscope (Thermo Fisher Scientific) operated at 300 kV configured for fringe-free illumination and equipped with a K2 direct electron detector (Gatan) mounted post Quantum 968 LS imaging filter (Gatan). The microscope was operated in EFTEM mode with a slit-width of 20 eV and using a 100 μm objective aperture. Automated data acquisition was performed using EPU (Thermo Fisher Scientific) and all images were collected using the K2 in counting mode. Ten-second movies were collected at a magnification of 165,000x and a pixel size of 0.84 Å, with a total dose of 64.8 e^-^/Å^2^ distributed uniformly over 40 frames. In total, 2437 movies were acquired with a realized defocus range of -0.5 to -2.5 μm.

CryoEM data analysis was performed in cryoSPARC version 3.2^50^ (**Extended Data Fig. 2, Supplementary Table 2**). Movies were motion-corrected using patch motion correction (multi) and CTF-estimated using patch CTF estimation (multi)^51^, and a 200-image subset was used for initial particle picking using the blob picker. Initial picks were subjected to 2D classification, and two classes were picked as templates for template-based particle picking of the entire 2437-image dataset. 2D classification of the resulting ∼2.5M particles revealed relatively few high-quality classes, likely a result of particle shape irregularity and high density on the grids. A ∼1M particle subset was used for initial ab initio 3D reconstruction, then the entire ∼2.5M particle dataset was used for heterogeneous refinement against these models, resulting in a 663,199-particle set that gave a ∼5.5 Å resolution map with recognizable protein features. This particle set was subjected to a further round of ab initio reconstruction and heterogeneous refinement to separate 2:2 Cap2:cGAS complexes from 2:1 complexes. The two separate particle sets were cleaned with another round of 3D classifications, then re-extracted with a 440-pixel (370 Å) box size and refined using the Non-Uniform Refinement NEW job type in cryoSPARC with the following options enabled: Maximize over-particle scale; Optimize per-particle defocus; Optimize per-group CTF params. For the 2:2 complex, C2 symmetry was applied during refinement. The resulting reconstructions showed resolution values of 2.74 Å (2:2 complex) and 2.91 Å (2:1 complex) using the 0.143-cutoff criterion of the Fourier shell correlations between masked independently refined half-maps. Resolution anisotropy for both reconstructions was assessed using the 3DFSC web server^52^.

An initial model for *E. cloacae* Cap2 was generated by AlphaFold2^53^. This model and the crystal structure of ATP-bound *E. cloacae* cGAS (PDB ID 7LJL^24^) were manually docked into the final 2:2 complex cryoEM map using UCSF Chimera^54^ and rebuilt in COOT^55^. For the E1 domain of Cap2 and for cGAS, high-resolution crystal structures were used to verify the accuracy of the resulting model. The final rebuilt model was real-space refined in phenix.refine^56^. This model was then docked into the 2:1 complex map, disordered regions were deleted, and the final model was real-space refined in phenix.refine. Structure validation was performed with MoProbity^57^ and EMRinger^58^. Structures were visualized in ChimeraX^54^ and PyMOL (Schrödinger, LLC).

### Crystallography

To determine a crystal structure of the Cap2 E1 domain bound to the cGAS C-terminus in the apo state, we cloned and purified a fusion construct with *E. cloacae* Cap2 residues 374-600 (C548A mutant) fused at its C-terminus to a flexible linker and residues 370-381 of cGAS (sequence: GSGKPAEPQKTGRFA). Purified protein was exchanged into a buffer containing 25 mM Tris-HCl pH 8.5, 200 mM NaCl, 5 mM MgCl_2_ and 1 mM TCEP, then concentrated to 30 mg/mL. Small rod-shaped crystals grew in hanging drop format by mixing 1:1 of protein with well solution containing 0.1 M Tris-HCl pH 8.5, 0.8 M LiCl, and 25% PEG 3350. Crystals were transferred to a cryoprotectant containing an additional 10% glycerol, then flash-frozen in liquid nitrogen. We collected a 1.77 Å resolution diffraction dataset at NE-CAT beamline 24ID-C at the Advanced Photon Source at Argonne National Laboratory (**Supplementary Table 4**). Data were processed with the RAPD pipeline, which uses XDS^59^ for data indexing and reduction, AIMLESS^60^ for scaling, and TRUNCATE^61^ for conversion to structure factors. We determined the structure by molecular replacement in PHASER^62^ using the refined Cap2 E1 domain structure from our cryoEM model of Cap2-cGAS. The model was rebuilt in COOT^55^, followed by refinement in phenix.refine^63^ using positional, individual B-factor, and TLS refinement (statistics in **Supplementary Table 4**).

To determine a crystal structure of the Cap2 E1 domain bound to the cGAS C-terminus in the AMP-bound reactive intermediate state, we cloned and purified a fusion construct with *E. cloacae* Cap2 residues 363-600 (C548A mutant) fused at its C-terminus to a flexible linker and residues 370-381 of cGAS (sequence: GSGKPAEPQKTGRFA). Purified protein was exchanged into crystallization buffer and concentrated to 30 mg/mL. A final concentration of 2.5 mM ATP was added to the protein and incubated overnight at 4 °C. Needle crystals grew in hanging drop format by mixing 1:1 of protein with well solution containing 0.1 M Tris-HCl pH 8.5, 0.2 M MgCl_2_, and 30% PEG 3350). Crystals were looped directly from the drop and flash-frozen in liquid nitrogen. A 2.11 Å resolution diffraction dataset was collected at NE-CAT beamline 24ID-E at Advanced Photon Source at Argonne National Laboratory and processed as above.

### Cap2/3 Biochemical Assays

For Cap2 activity assays, the indicated combinations of *E. cloacae* His_6_-Cap2 and untagged cGAS (wild-type or mutant) were coexpressed in Rosetta2 pLys *E. coli* cells, then purified as above using a Ni^2+^ affinity column. Samples were analyzed by SDS-PAGE with Coomassie staining. For quantitation, experiments were run in triplicate and Coomassie blue-stained bands quantified using Fiji software^64^. For Cap3 activity assays, model substrates comprising *E. cloacae* or *V. cholerae* His_6_-cGAS (wild-type or mutant) fused at their C-terminus to GFP were cloned and purified as above. Model substrates (4.5 μg) were incubated with Cap3 (1.5 μg) in a reaction buffer with 20 mM HEPES pH 7.5, 100 mM NaCl, 20 mM MgCl_2_, 20 μM ZnCl_2_, and 1 mM DTT (20 μL total reaction volume). Reactions were incubated 30 minutes at 37 °C, then analyzed by SDS-PAGE with Coomassie blue staining.

### Trypsin Mass Spectrometry

For trypsin mass spectrometry of purified proteins (HA-cGAS and Cap2-GFP), in-gel digestion was performed according to a previously described method^65^. Briefly, proteins in diced gel bands were reduced by 100 μL of 10 mM DTT (Dithiothreitol) for 30 minutes at 37 °C and then alkylated by 6 μL of 0.5 M iodoacetamide in water for 20 minutes at room temperature in the dark. To digest proteins, 25-30 μL of 10 ng/μL Trypsin (Promega, V511A) in 50 mM ammonium bicarbonate (pH 8) was added to cover the gel pieces and incubated on ice for 30 minutes until fully swollen. An additional 10-20 μL of ammonium bicarbonate buffer was added and the sample was incubated overnight at 37 °C. The next day, trypsin digested peptides were extracted from the gel via multiple solvent extractions, dried under vacuum and then resuspended in 5 μL of 0.6% acetic acid. The digested peptides were analyzed by a Thermo Fisher Scientific Orbitrap Fusion LUMOS Tribrid mass spectrometer using standard LC-MS/MS method^65^.

MS data analysis was performed using the Trans-Proteomic Pipeline (TPP, Seattle Proteome Center). Briefly, MS data were searched using the search engine COMET against a composite *E. coli* database that additionally contained protein sequences for *E. cloacae* cGAS and Cap2, plus common contaminants. Variable modifications include possible oxidation of methionine (15.9949 Da) and expected FA remnant of the cGAS C-terminus (218.10552 Da); and a static modification of cysteine by IAA (57.021464 Da) was included. The COMET search results were further analyzed with PeptideProphet and ProteinProphet^66^. Peptides with a probability of >0.9 and mass accuracy of <10 ppm were subjected to further manual inspection of the MS/MS spectra to confirm major fragment ions are accounted for.

---

**Extended Data Figure 1.**
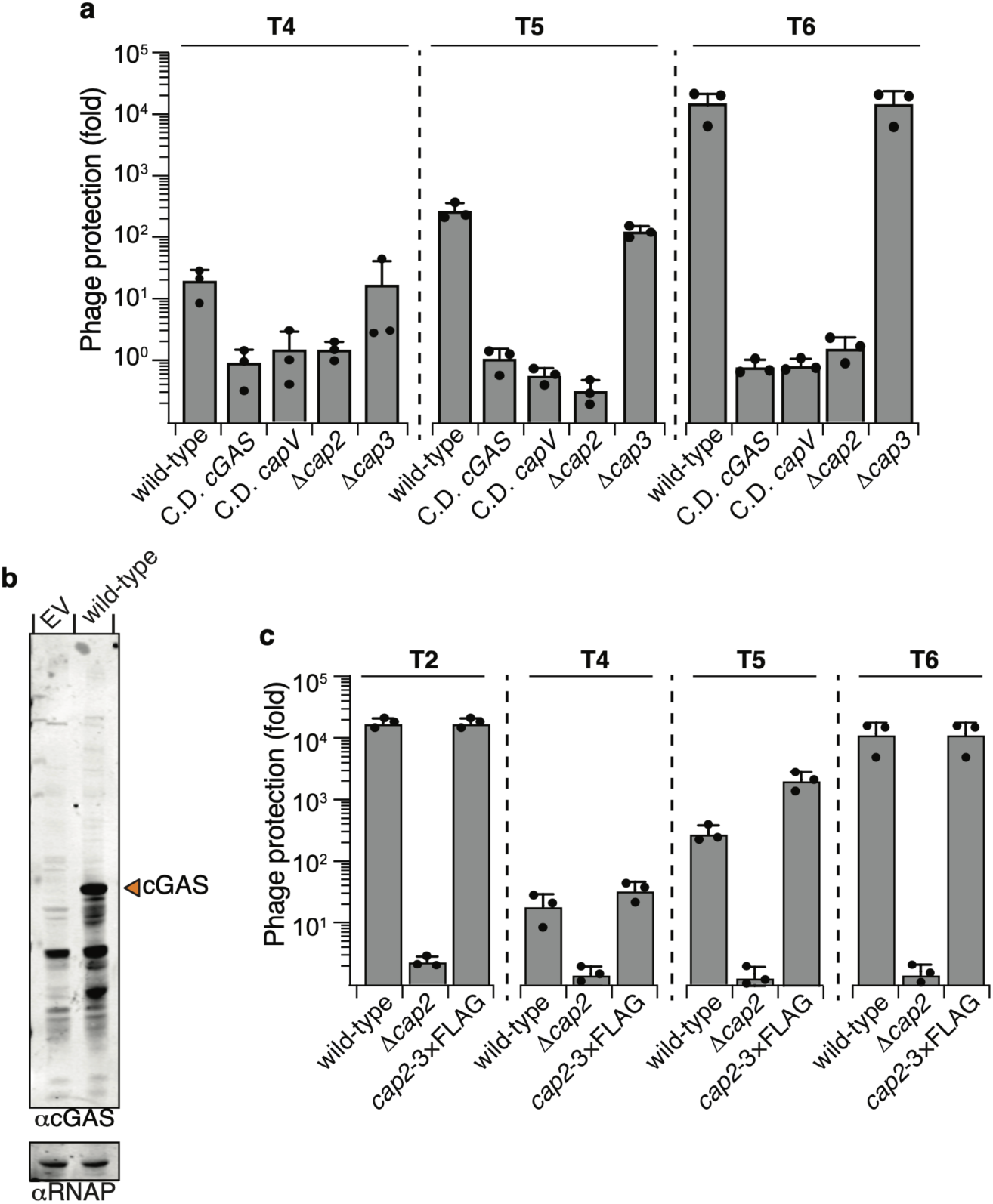
Phage protection assays and cGAS antibody verification. **(a)** Efficiency of plating of the indicated phage when infecting *E. coli* expressing CBASS with the indicated genotype. Data plotted as in Fig. 1b. C.D. cGAS: DID131AIA.; C.D. *capV*: C62A. **(b)** Whole cell western blot analysis of *E. coli* expressing either an empty vector (EV) or CBASS (wild-type). αcGAS western blot used a custom cGAS antibody; orange arrow indicates monomeric cGAS at the expected molecular weight. αRNAP western blot serves as a loading control for bacterial cells. **(c)** Efficiency of plating of the indicated phage when infecting *E. coli* expressing *V. cholerae* CBASS with the indicated genotype. Data plotted as in **Fig. 1b**.

**Extended Data Figure 2.**
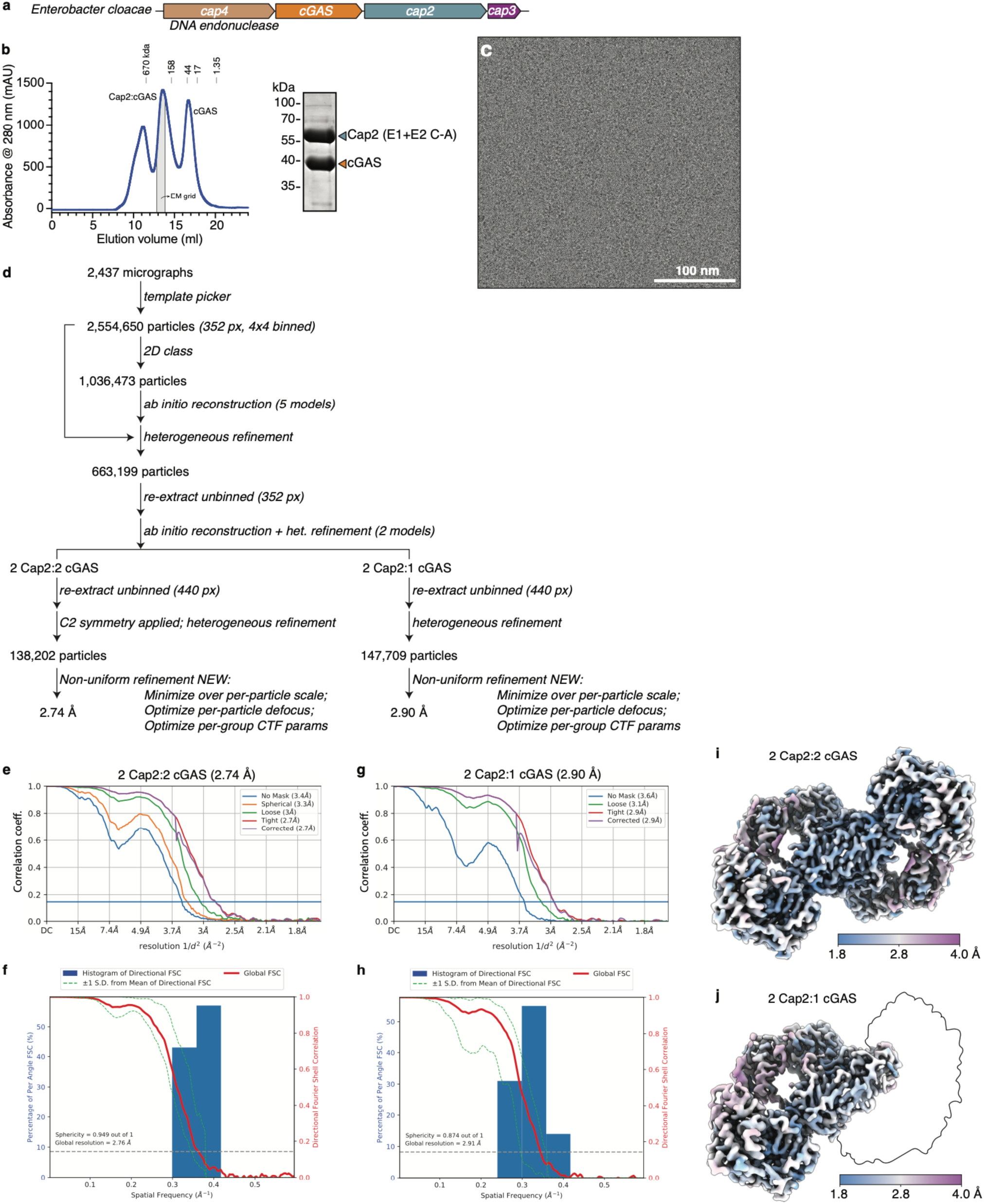
CryoEM Workflow. **(a)** Operon structure of CBASS from *E. cloacae*; see **Supplementary Table 7** for relevant accession numbers. **(b)** Size exclusion chromatography elution profile (Superdex 200 Increase 10/300 GL) and SDS-PAGE analysis of *E. cloacae* Cap2–cGAS. The fraction used for cryoEM analysis is shaded in gray. **(c)** Representative electron micrograph of *E. cloacae* Cap2–cGAS. **(d)** Workflow for structure determination in cryoSPARC^50^. Templates used for particle picking were chosen from initial 2D classes from blob-picked particles in a 200-image subset. **(e)** Fourier Shell Correlation (FSC) curve for the final refinement of the 2:2 Cap2–cGAS complex. **(f)** 3D FSC analysis^52^ for the 2:2 Cap2–cGAS complex. **(g)** Fourier Shell Correlation (FSC) curve for the final refinement of the 2:1 Cap2–cGAS complex. **(h)** 3D FSC analysis for the 2:1 Cap2–cGAS complex. **(i)** Local resolution of the final refined map for the 2:2 Cap2–cGAS complex, colored from blue (1.8 Å or higher) to magenta (4.00 Å or lower). **(j)** Local resolution of the final refined map for the 2:1 Cap2–cGAS complex colored from blue (1.8 Å or higher) to magenta (4.00 Å or lower). Outline indicates the areas of missing density compared to the 2:2 Cap2–cGAS complex.

**Extended Data Figure 3.**
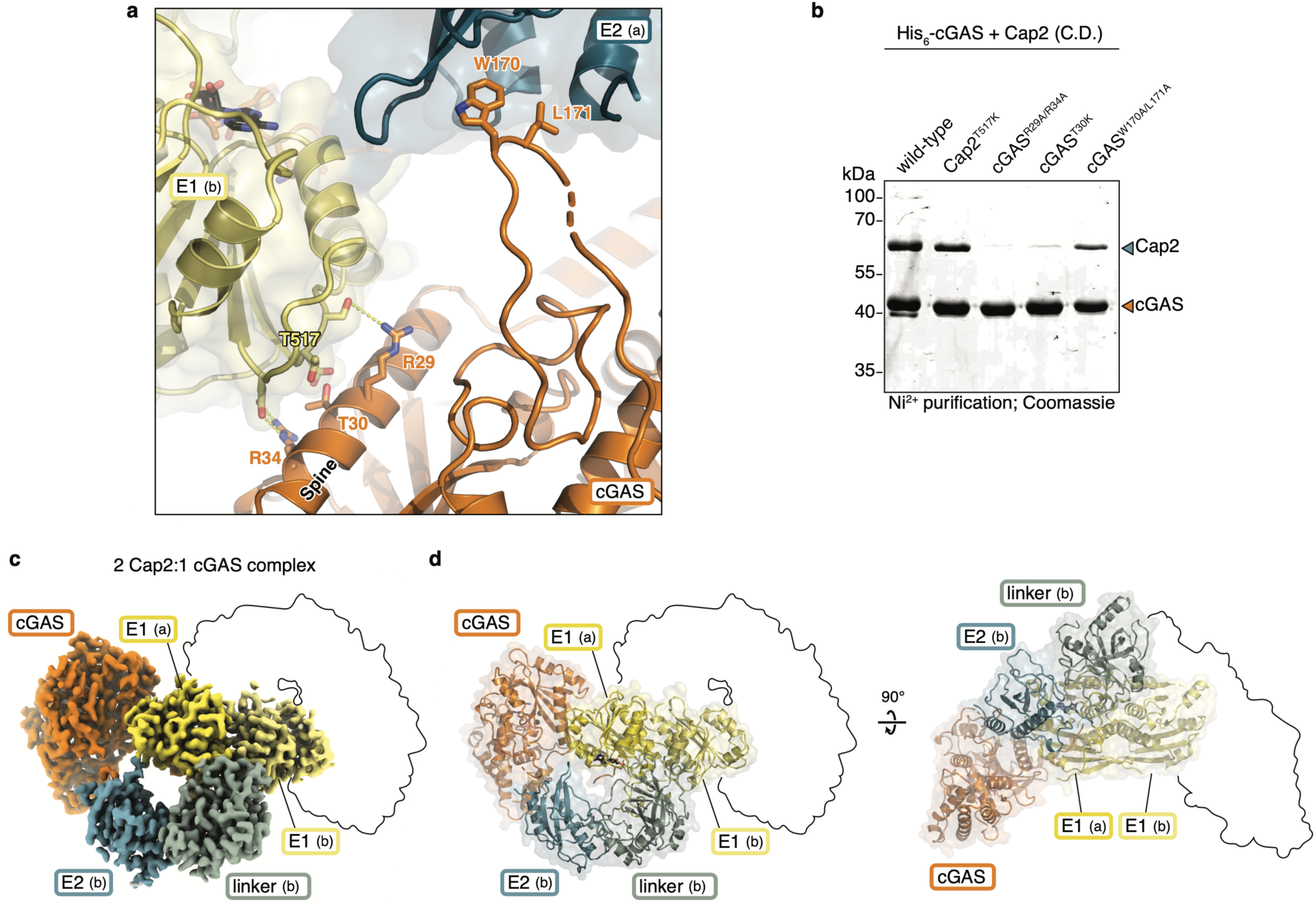
cGAS rigidifies Cap2 through a bipartite interaction. **(a)** Closeup view of the interaction between Cap2 (yellow/blue) and cGAS (orange), with key residues shown as sticks and labeled. **(b)** Coomassie stained SDS-PAGE of proteins that purified by Ni^2+^-affinity chromatography from *E. coli* co-expressing *E. cloacae* 6×His-tagged *cGAS* (His_6_-cGAS) and catalytically inactivated *cap2* (C548A/C109A; C.D.) with the indicated genotype. **(c)** CryoEM density for 2:1 Cap2–cGAS complex, with domains labeled and colored as in **Figure 2b**. Outline indicates the areas of missing density compared to the 2:2 Cap2–cGAS complex, including one protomer of cGAS and the E2 and linker domains for the (a) protomer of Cap2. **(d)** Two views of the 2:1 Cap2–cGAS complex, with domains labeled and colored as in panel (c). Outlines indicate the areas of missing density compared to the 2:2 Cap2–cGAS complex.

**Extended Data Figure 4.**
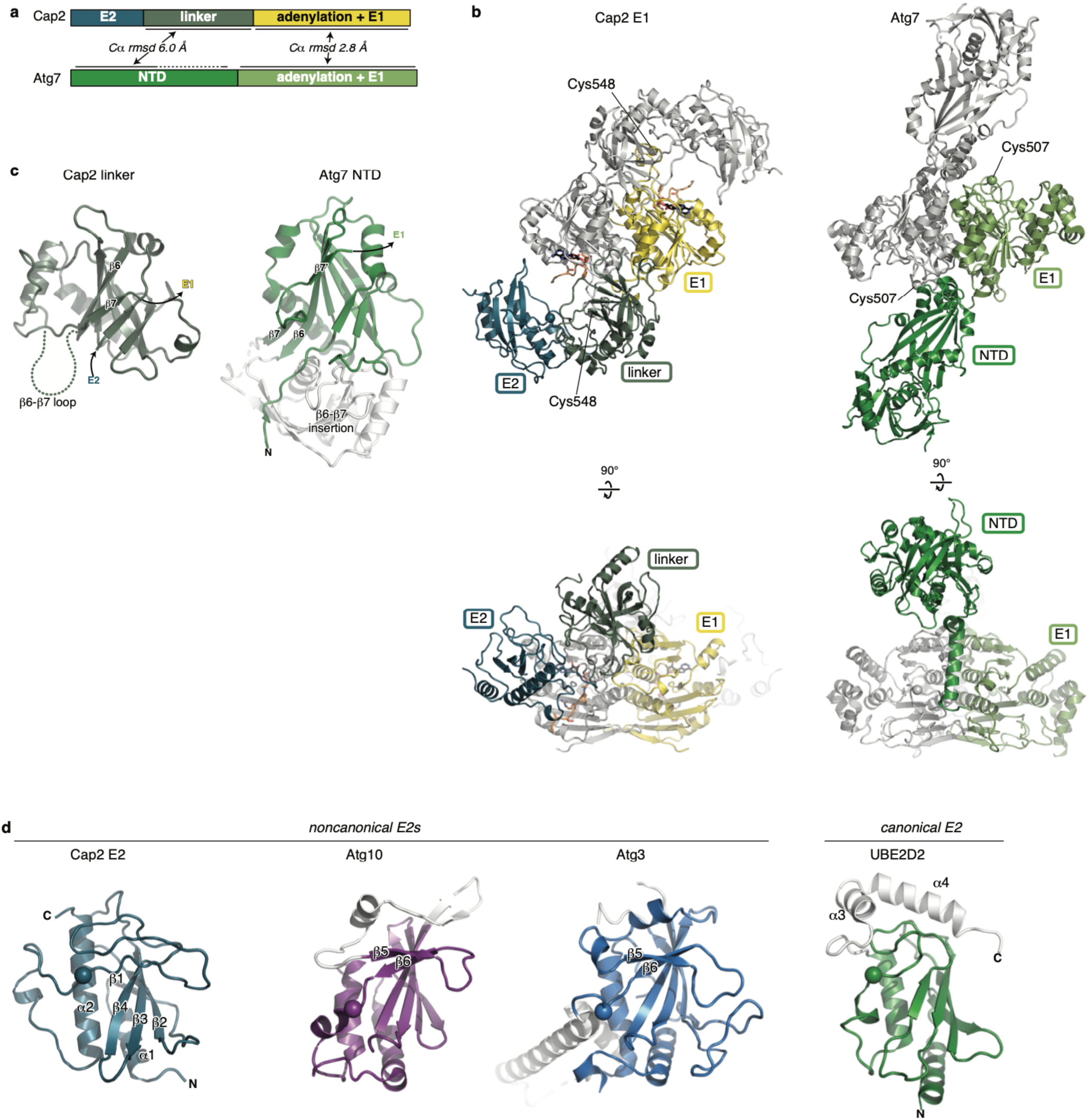
Cap2 is related to autophagy E1 and E2 proteins. **(a)** Domain schematic of *E. cloacae* Cap2 and *S. cerevisiae* ATG7, with approximate root-mean-squared distance (Cα r.m.s.d.) values for the linker/NTD and E1 domains noted. **(b)** Structures of *E. cloacae* Cap2 (left), compared to *S. cerevisiae* ATG7 (right; PDB ID 4GSK^28^), with one protomer colored as in panel (a) and the dimer mate colored gray. For each protein, the E1 active-site cysteine residue (C548 for Cap2, C507 for ATG7) is shown as a sphere and labeled. **(c)** Structures of the *E. cloacae* Cap2 linker domain (left), compared to the *S. cerevisiae* ATG7 NTD (right; PDB ID 4GSK^28^). ATG7 features a second subdomain (residues 147-268, shown in white) inserted into the loop separating β-strands 6 and 7 (labeled) where Cap2 has a partially disordered loop (residues 319-356). **(d)** Structure of the Cap2 E2 domain (active-site C109 shown as a sphere), compared to *Kluyveromyces marxianus* ATG10 (PDB ID 3VX7^29^), *S. cerevisiae* ATG3 (PDB ID 2DYT^73^), and *Homo sapiens* UBE2D2 (PDB ID 4DDG^74^). Structural features not shared are shown in white. The active-site cysteine of each protein is shown as a sphere.

**Extended Data Figure 5.**
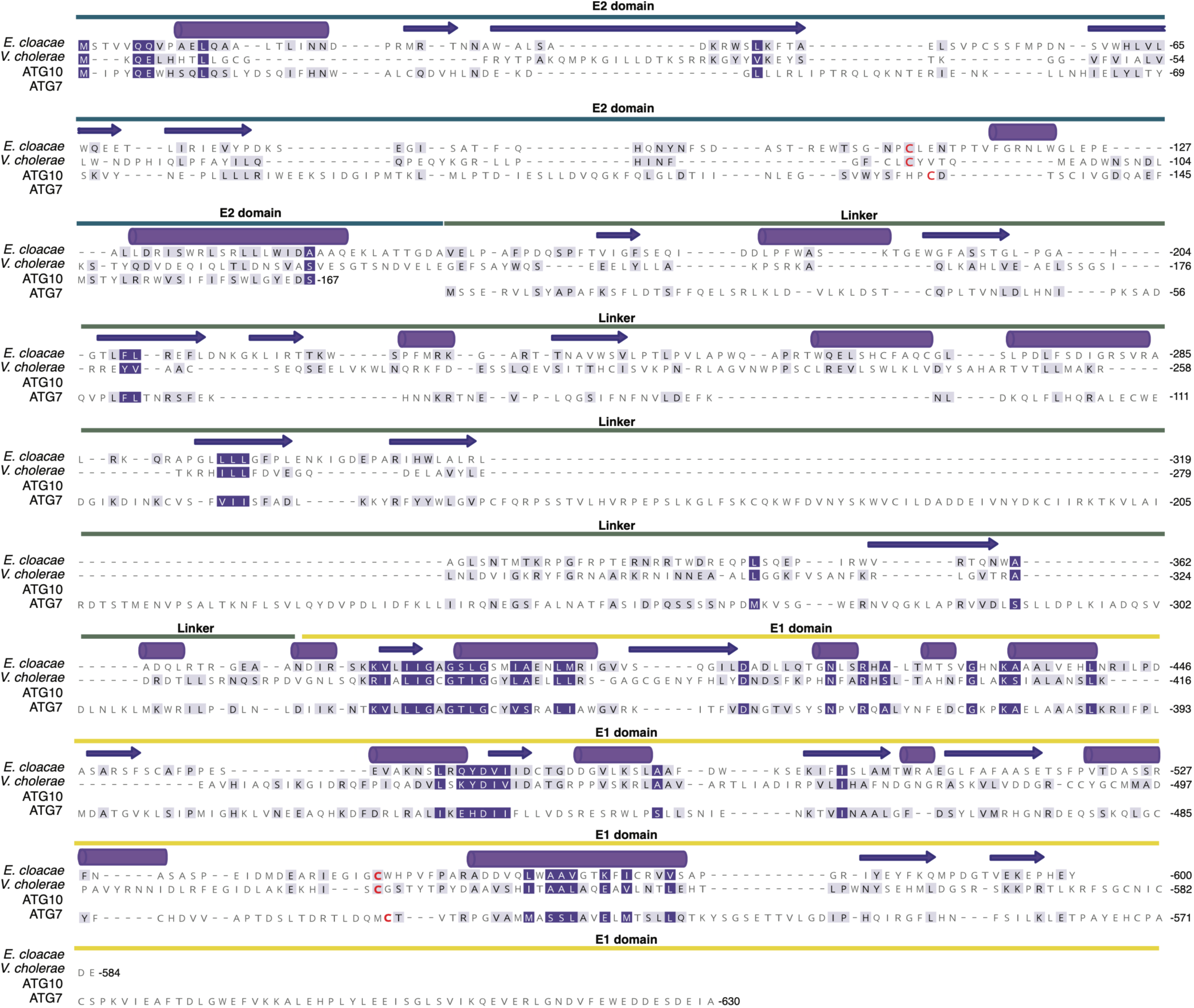
Cap2 protein alignment. **(a)** Protein alignment of Cap2 from *E. cloacae*, Cap2 from *V. cholerae*, ATG10 from *S. cerevisiae* (4EBR^75^), and ATG7 from *S. cerevisiae* (3T7H^25^). Domains are indicated above the alignment with colors corresponding to **Fig. 2**. The secondary structure of Cap2 from *E. cloacae* is indicated in purple with alpha helices depicted as cylinders and beta sheets as arrows. The catalytic cysteines found in the E2 and E1 domains are highlighted in red.

**Extended Data Figure 6.**
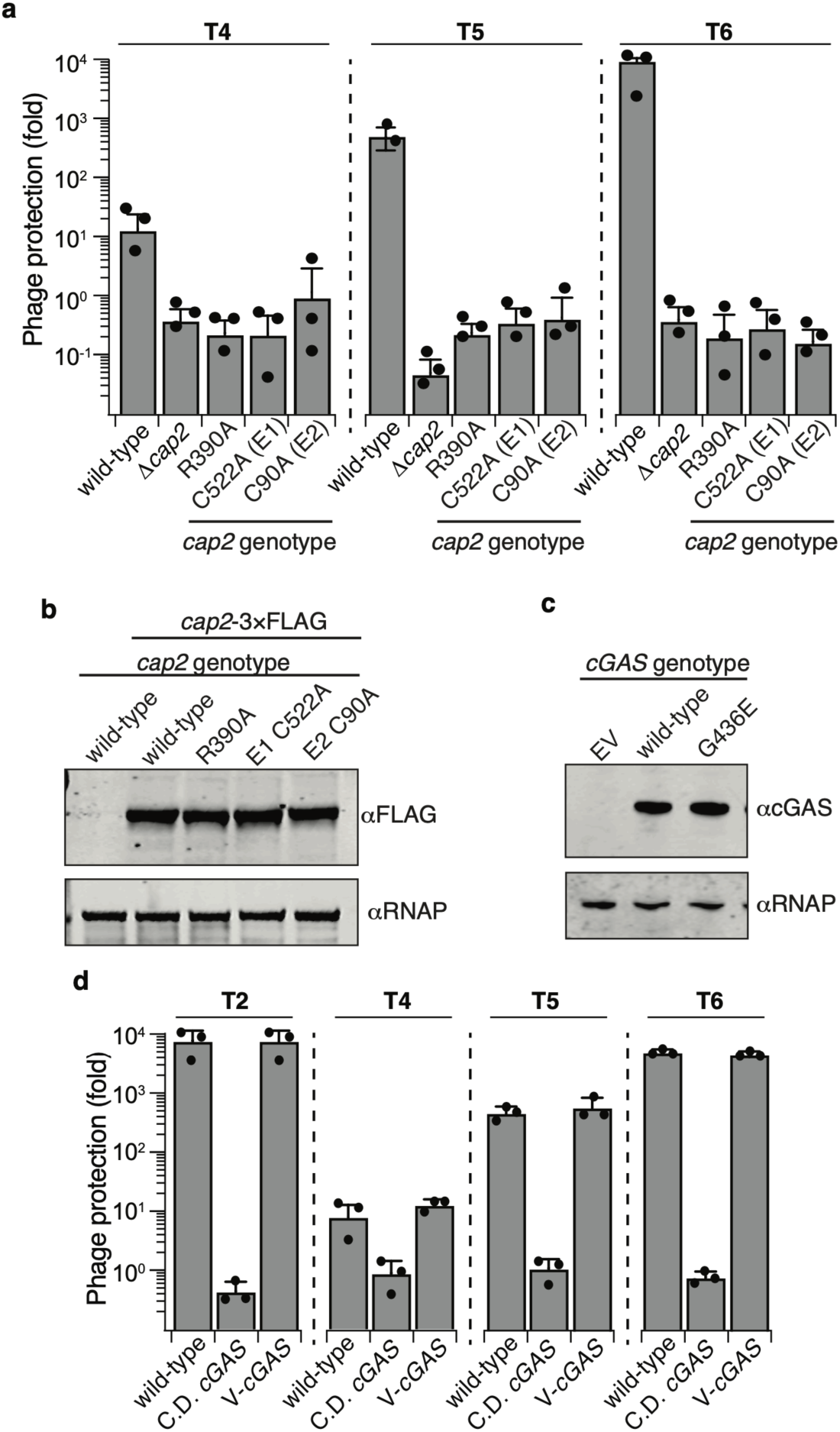
Phage protection and western blot analysis of Cap2 mutants and tagged cGAS. **(a)** Efficiency of plating of the indicated phage when infecting *E. coli* expressing CBASS with the indicated genotype. Data plotted as in **Fig. 1b. (b)** Western blot analysis of cell lysates from *E. coli* expressing CBASS with the indicated genotypes demonstrating that the mutations do not affect expression levels. **(c)** Western blot analysis of cell lysates from *E. coli* expressing CBASS with the indicated genotypes demonstrating that the mutations do not affect protein expression levels. **(d)** Efficiency of plating of the indicated phage when infecting *E. coli* expressing CBASS with the indicated genotypes. Data plotted as in **Fig. 1b**. V-cGAS: N-terminally VSV-G epitope tagged cGAS.

**Extended Data Figure 7.**
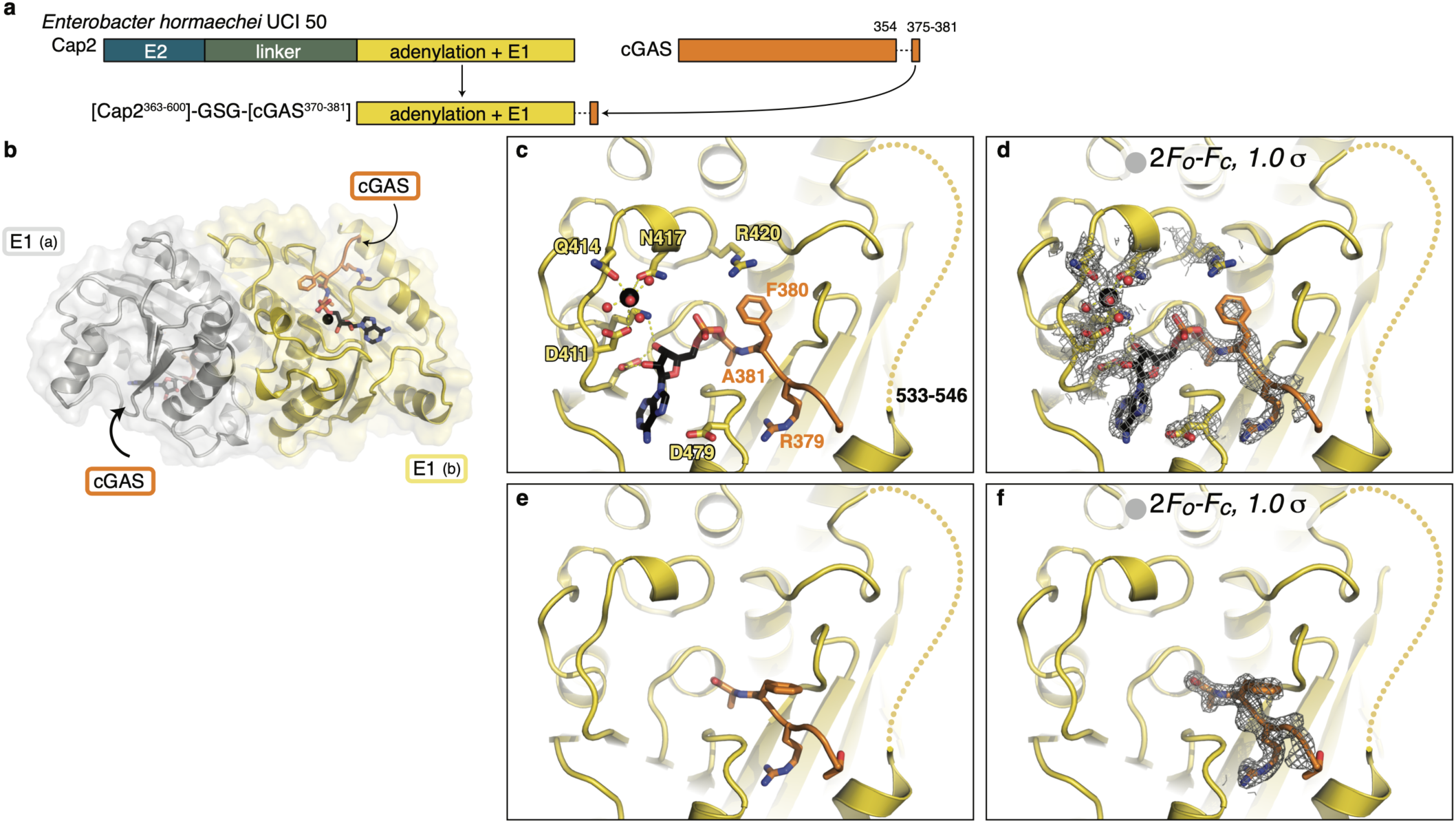
Crystal structures of an *E. cloacae* Cap2 E1-cGAS fusion. **(a)** Design of a fusion between the C-terminal E1 domain of *E. cloacae* Cap2 (residues 363-600) and the C-terminus of cGAS (residues 370-381), with a three-residue GSG linker. **(b)** 2.1 Å resolution crystal structure of the *E. cloacae* Cap2–cGAS fusion crystallized in the presence of ATP, with two Cap2 E1 domains colored yellow and gray, and the two cGAS C-termini colored orange. See also **Supplementary Table 4. (c)** Closeup of the Cap2 adenylation active site, showing the cGAS–AMP conjugate and active site residues. Residues 533-546 are disordered and represented by a dotted line. Bound Mg^2+^ ion is shown in black. **(d)** View as in (c), with 2*F*_*O*_*-F*_*C*_ electron density contoured at 1.0 σ around the cGAS–AMP conjugate and active site residues. **(e)** Closeup of the Cap2 adenylation active site in a 1.8 Å-resolution structure of the Cap2-cGAS fusion crystallized in the absence of added nucleotide (apo state). **(f)** View as in (e), with 2*Fo-Fc* electron density contoured at 1.0 σ around the cGAS C-terminus.

**Extended Data Figure 8.**
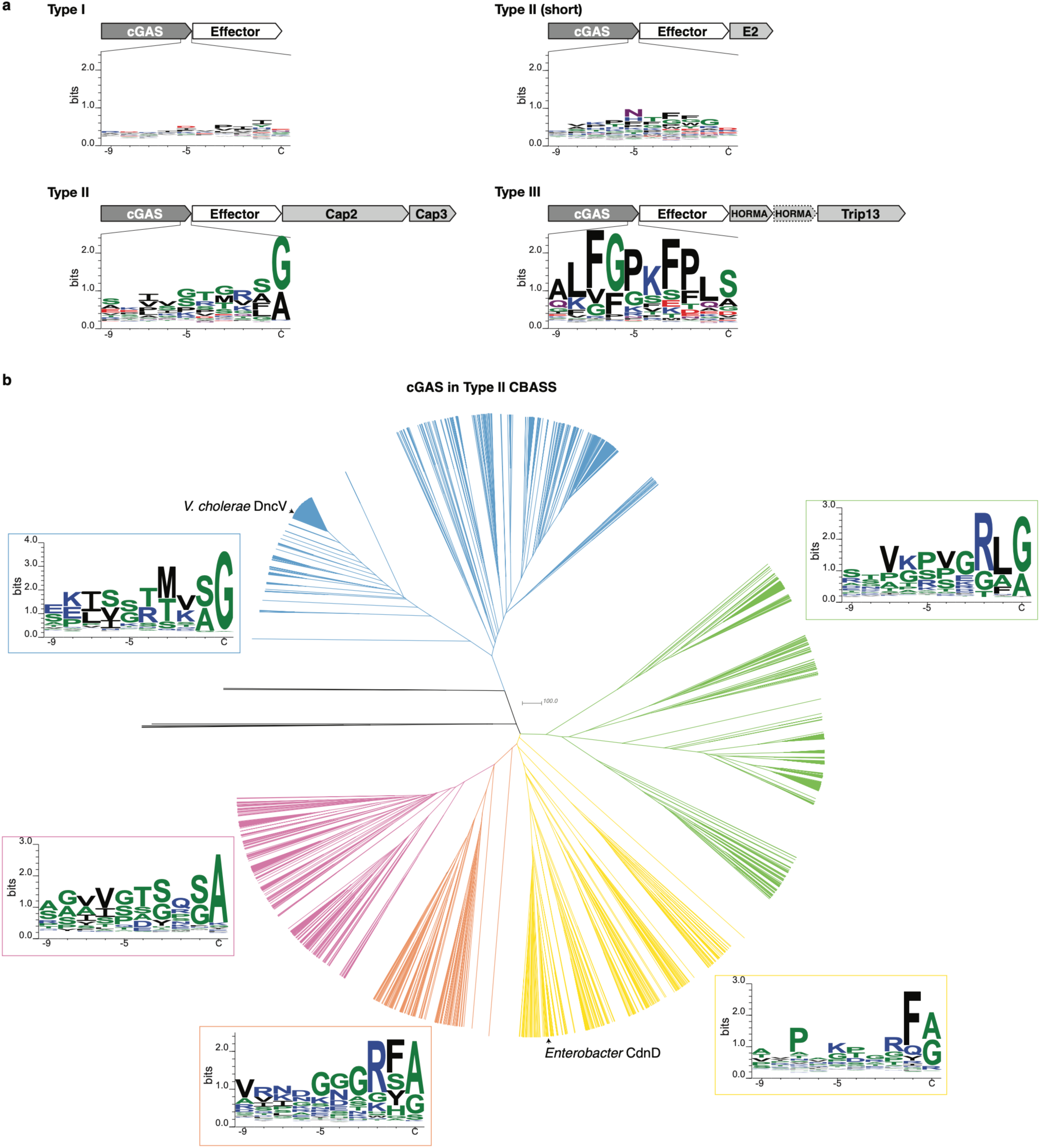
The C-terminus of cGAS is conserved in Type II CBASS systems. **(a)** Sequence logos of C-terminal 10 residues of cGAS in Type I (2284 sequences), Type II (1556 sequences), Type II (short) (593 sequences), and Type III (540 sequences) CBASS systems^7^. Type II (short) CBASS systems encode an E2 ubiquitin transferase-like enzyme without a linked E1 domain, and do not encode a JAB isopeptidase. **(b)** Phylogenetic tree of 2284 cGAS enzymes in Type II CBASS systems^7^, with C-terminus sequence logos for each major branch. *V. cholerae* cGAS (DncV) and *E. cloacae* cGAS (CdnD) are labeled.

**Extended Data Figure 9.**
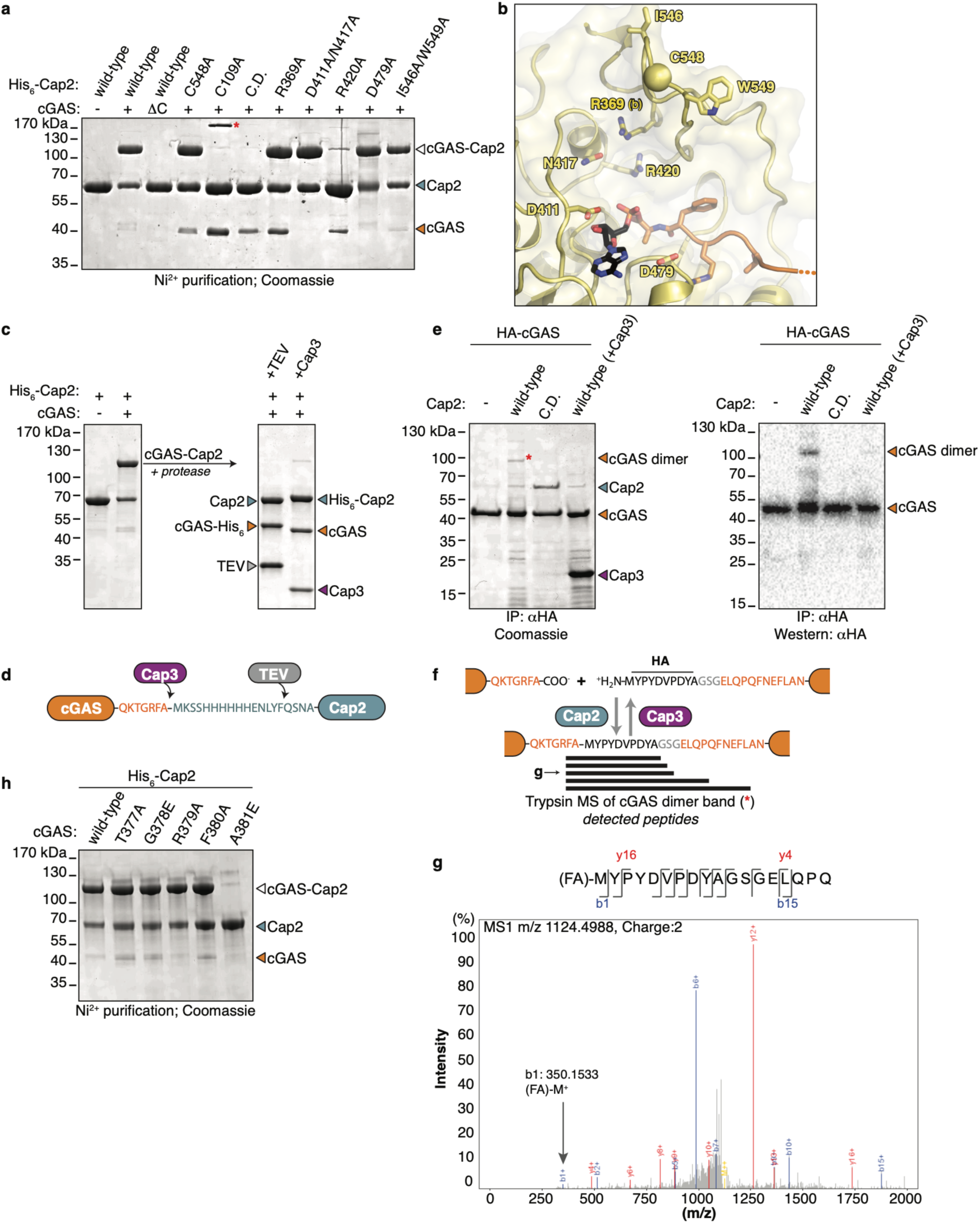
Cap2 conjugates the cGAS C-terminus to a target. **(a)** SDS-PAGE analysis of *E. cloacae* Cap2 activity assay. The indicated genotypes of His_6_-Cap2 and cGAS were expressed from a single plasmid and the formation of a cGAS–His_6_-Cap2 conjugate was used as an indicator of Cap2 activity. (-): no cGAS; (+): wild-type cGAS; (ΔC): cGAS lacking its C-terminus (19 residues deleted); C.D.: C548A/C109A catalytically dead Cap2. Red asterisk: putative intermediate with cGAS thioester-linked to the Cap2 E1 catalytic cysteine (C548). The formation of a cGAS–Cap2 conjugate in the absence of a functional E1 catalytic cysteine (C548A) indicates that *in vitro*, this residue is dispensable for catalysis and the nearby E2 catalytic cysteine (C109) can function in its stead. **(b)** Cap2 E1 active site (yellow) in Cap2–cGAS cryoEM structure with the residues mutated in (a) indicated and the E1 active-site cysteine residue (C548 for Cap2) shown as a sphere and labeled. cGAS C-terminus (orange) conjugated to AMP (black). **(c)** *Left:* SDS-PAGE analysis of Ni^2+^-purified *E. cloacae* His_6_-Cap2, expressed either alone or with full-length cGAS. Right: Protease treatment (TEV or Cap3) of the cGAS–His_6_-Cap2 conjugate. **(d)** Schematic of the inferred cGAS–His_6_-Cap2 conjugate formed upon coexpression of *E. cloacae* His_6_-Cap2 and cGAS, with cleavage sites for Cap3 and TEV protease indicated. **(e)** SDS-PAGE analysis with detection by Coomassie (left) or αHA Western blot (right) of αHA immunopurified *E. cloacae* Cap2 coexpressed with HA-cGAS. C.D.: Cap2 C548A/C109A. Red asterisk indicates band used for tryptic mass spectrometry analysis in (f-g). **(f)** Peptides detected in tryptic mass spectrometry of the marked band in (e), showing conjugation of cGAS to the N-terminus of a second HA-cGAS molecule. See **Supplementary Table 9** for mass spectrometry data. **(g)** Collision-induced dissociation mass spectrum of the peptide indicated in (f), with b1 peak indicated (mass of 350.1533 is that of Met+(H+)+(Phe-Ala)). **(h)** SDS-PAGE analysis of Ni^2+^-purified *E. cloacae* His_6_-Cap2 with cGAS with the indicated genotype.

**Extended Data Figure 10.**
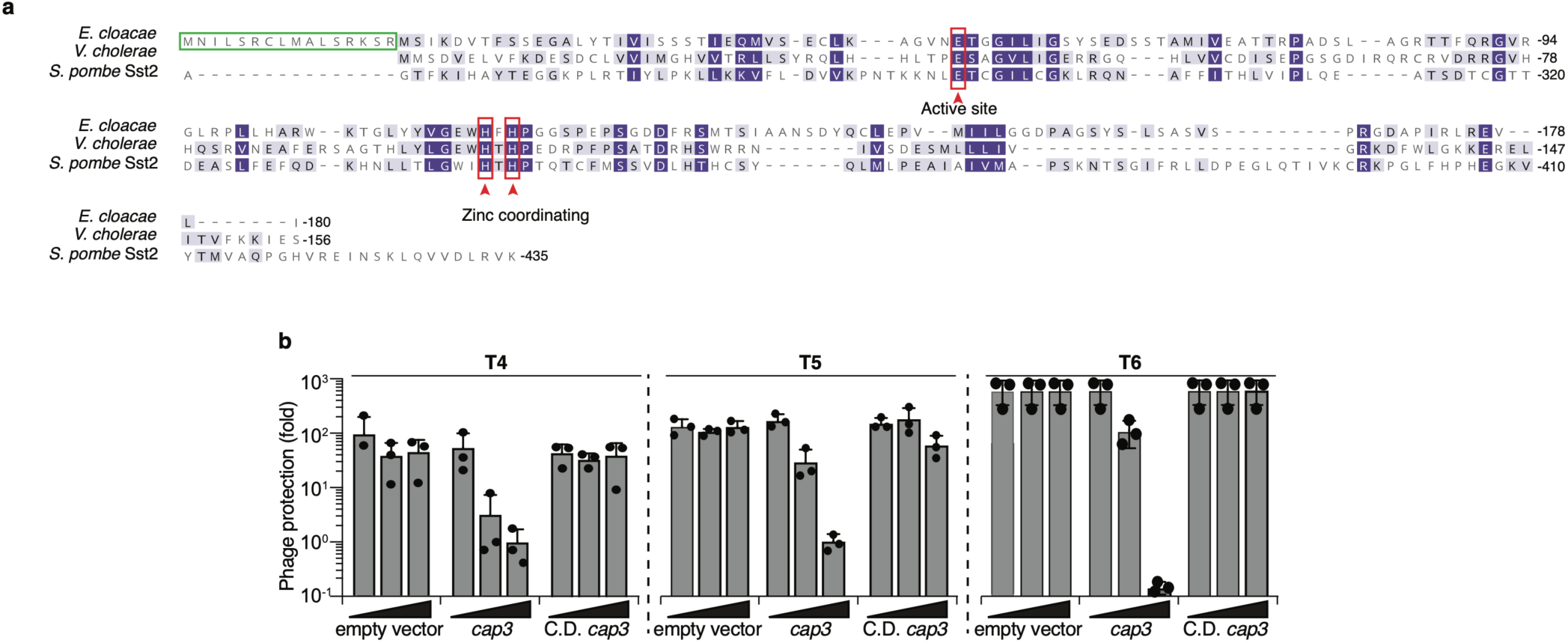
Cap3 overexpression inhibits phage protection. **(a)** Protein alignment of the JAMM/JAB protease Sst2 from *S. pombe* (Uniprot ID Q9P371; residues 235-435), Cap3 from *E. cloacae*, and Cap3 from *V. cholerae*. The active site glutamate, as well as two zinc-coordinating histidine residues, are noted. For experiments using Cap3 from *E. cloacae*, the first 16 annotated amino acids (green box) were removed as we found they are unlikely to represent a true translation start site. **(b)** Efficiency of plating of the indicated phage infecting *E. coli* expressing CBASS Δ*cap3* in the absence or presence of overexpressed *cap3* with the indicated genotype. Data plotted as in **Fig. 1b**. C.D. *cap3*: HTH101ATA.

**Extended Data Figure 11.**
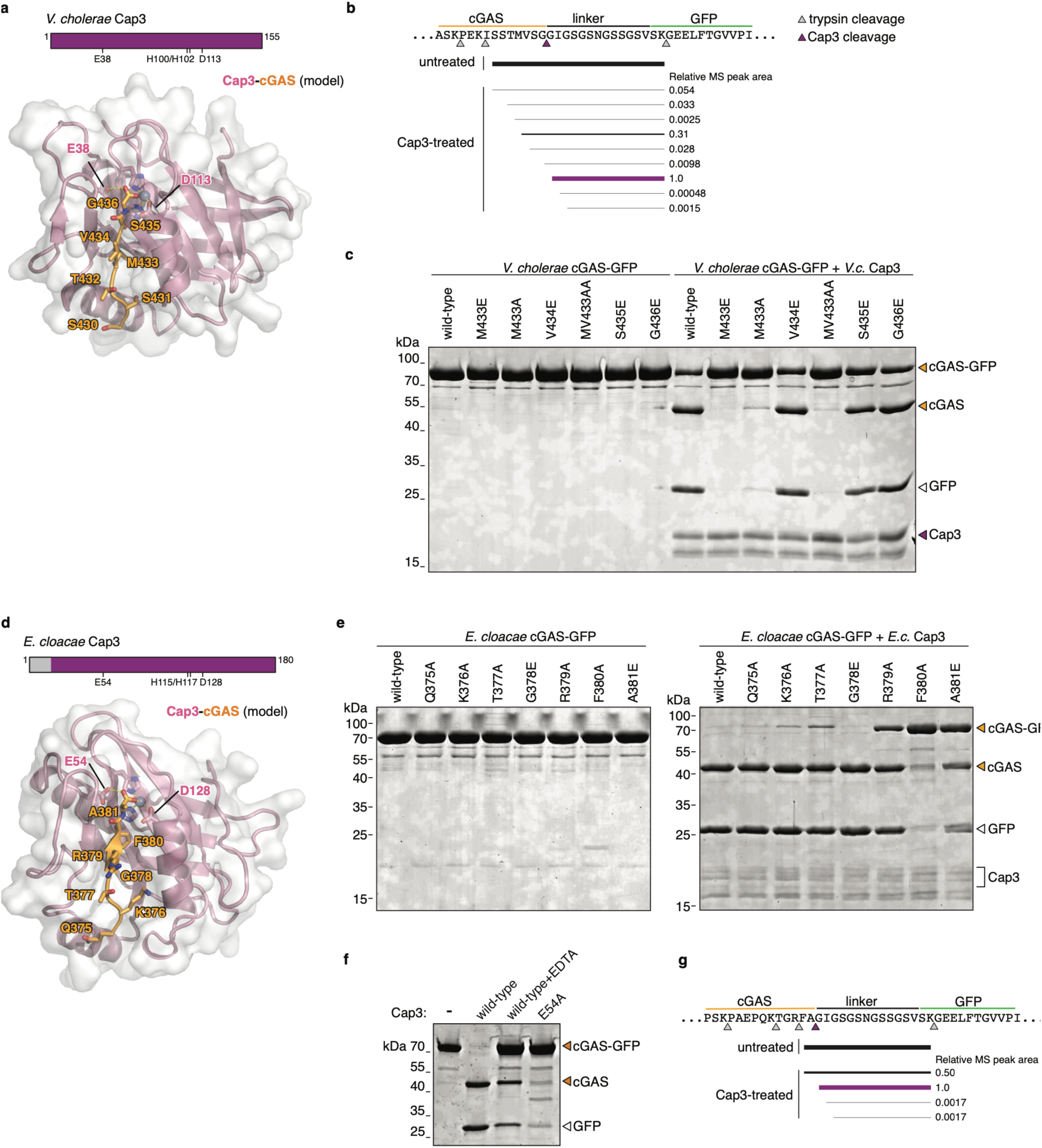
Cap3 cleavage of a cGASylated model substrate. **(a)** Domain schematic and model of the *V. cholerae* Cap3-cGAS complex, generated by AlphaFold2^53^ and with the cGAS C-terminus and Zn^2+^ ion manually modeled from an overlay with a structure of *S. pombe* Sst2 bound to ubiquitin^76^ (PDB ID 4K1R;). **(b)** Summary of tryptic digest mass spectrometry analysis of the *V. cholerae* Cap3-treated cGAS bands as in **Fig. 4b**. Pink arrow indicates the inferred Cap3 cleavage site; gray arrows indicate trypsin cleavage sites. See **Supplementary Table 5** for data. **(c)** Coomassie stained SDS-PAGE of a *V. cholerae* model substrate (cGAS-GFP fusion protein) with the indicated mutations in the cGAS C-terminus, with and without incubation with *V. cholerae* Cap3. **(d)** Domain schematic and model of the *E. cloacae* Cap3-cGAS complex, generated by AlphaFold2^53^ and with the cGAS C-terminus and Zn^2+^ ion manually modeled from an overlay with a structure of *S. pombe* Sst2 bound to ubiquitin^76^ (PDB ID 4K1R). **(e)** Coomassie stained SDS-PAGE of an *E. cloacae* model substrate (cGAS-GFP fusion protein) with the indicated mutations in the cGAS C-terminus, with and without incubation with *E. cloacae* Cap3. **(f)** Coomassie stained SDS-PAGE of an *E. cloacae* model substrate (cGAS-GFP fusion protein) incubated with *E. cloacae* Cap3 with the indicated reaction condition/genotype. **(g)** Summary of tryptic digest mass spectrometry analysis of the *E. cloacae* Cap3-treated cGAS bands as in (f), showing the putative Cap3 cleavage site. Pink arrow indicates the inferred Cap3 cleavage site; gray arrows indicate trypsin cleavage sites. See **Supplementary Table 5** for data.

**Supplementary Table S2.** CryoEM Data collection, reconstruction, and refinement statistics.

|  | Cap2-cGAS 2:2 complex | Cap2-cGAS 2:1 complex |
| --- | --- | --- |
| Data collection |  |  |
| Microscope | TFS Titan Krios G3 |  |
| Voltage (keV) | 300 |  |
| Nominal magnification | 165,000 |  |
| Spherical aberration (mm) | 2.7 |  |
| Detector | Gatan K2 |  |
| Pixel size (Å) | 0.8252 |  |
| GIF slit-width (eV) | 20 |  |
| Fluence (e-/Å<sup>2</sup>) | 65 |  |
| Defocus range (μm) | -0.5 to -2.5 |  |
| Movies collected | 2437 |  |
| Reconstruction |  |  |
| Particle number | 138,202 | 147,709 |
| Symmetry applied | C2 | none |
| Resolution (FSC<sub>0.143</sub>, Å) | 2.74 | 2.9 |
| Local resolution range, 25th- 75th percentile (FSC<sub>0.143</sub>, Å) | 2.58-5.47 | 2.71-5.70 |
| 3D FSC sphericity <sup>a</sup> | 0.949 | 0.874 |
| Refinement |  |  |
| Model Composition |  |  |
| Non-hydrogen atoms | 14764 | 8690 |
| Protein residues | 1850 | 1096 |
| Ligands | 4 Mg<sup>2+</sup>, 2 AMP, 2 ADP, 2 ATP | 2 Mg<sup>2+</sup>, 1 AMP, 1 ADP, 1 ATP |
| Isotropic ADPs (Å<sup>2</sup>) |  |  |
| Protein | 99.97 | 116.24 |
| Ligands | 90.14 | 110.7 |
| R.m.s deviations |  |  |
| Bond lengths (Å) | 0.002 | 0.003 |
| Bond angles (°) | 0.544 | 0.519 |
| Validation |  |  |
| Molprobability score | 1.11 | 1.14 |
| Molprobability clashscore | 3.2 | 3.15 |
| Rotamer outliers (%) | 0 | 0 |
| EMRinger score <sup>b</sup> | 3.77 | 3.99 |
| Ramachandran (%) |  |  |
| Favored | 98.14 | 97.87 |
| Allowed | 1.86 | 2.13 |
| Outliers | 0 | 0 |
| Map-Model FSC<sub>0.143</sub>/FSC<sub>0.5</sub> (Å) | 2.7/3.0 | 2.9/3.1 |
| PDB ID <sup>c</sup> | 7TO3 | 7TQD |
| EMDB ID <sup>d</sup> | 26028 | 26066 |

<sup>a</sup> 3D FSC values were calculated by the Remote 3DFSC Processing Server (https://3dfsc.salk.edu) (ref. 52).
<sup>b</sup> EMRinger scores were calculated by EMRinger (ref. 58).
<sup>c</sup> Coordinates have been deposited in the RCSB Protein Data Bank (http://www.rcsb.org).
<sup>d</sup> EM maps have been deposited with the Electron Microscopy Data Bank (https://www.ebi.ac.uk/emdb/).

**Supplementary Table S3.**
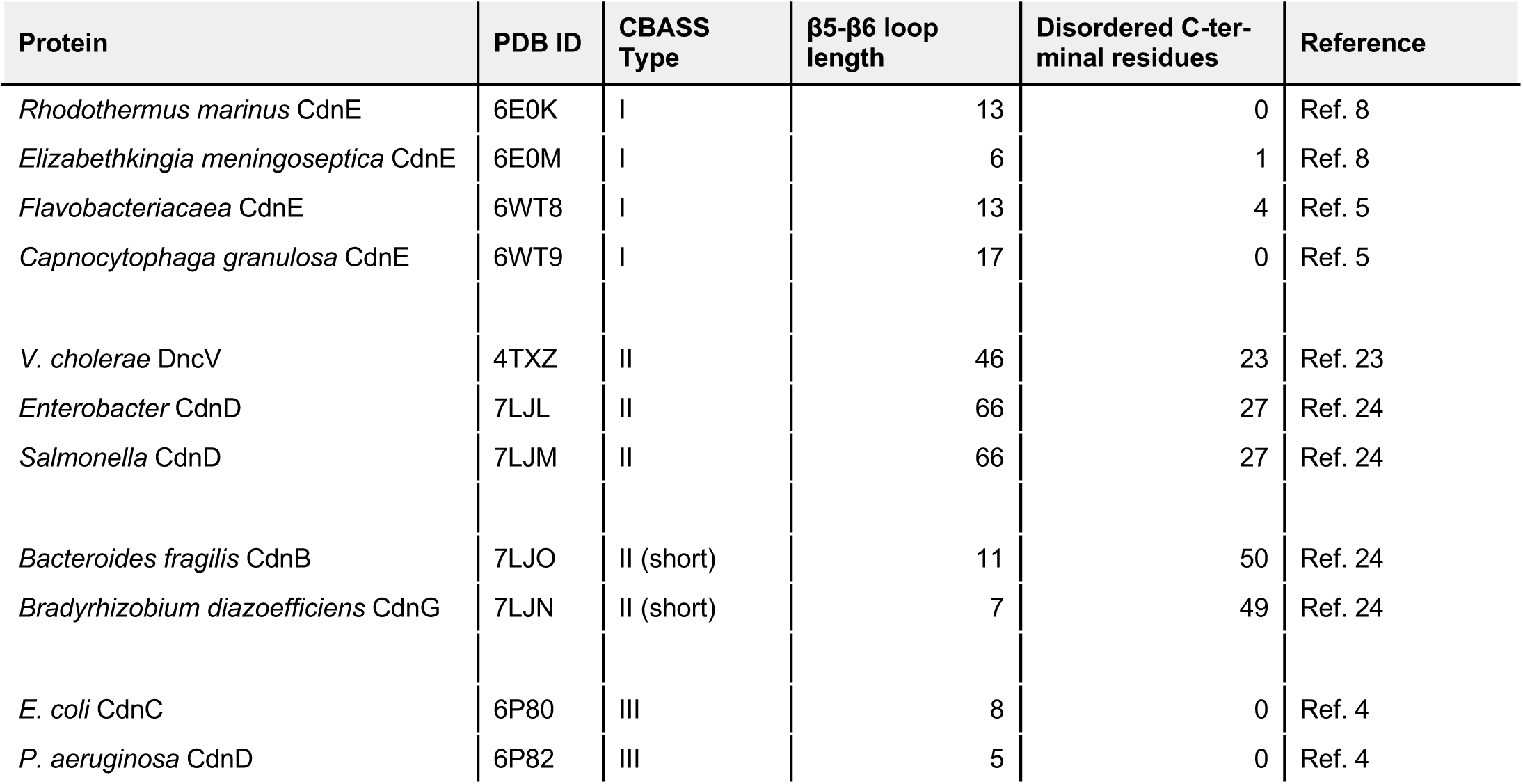
Structural features of bacterial cGAS proteins.

| Protein | PDB ID | CBASS Type | β5-β6 loop length | Disordered C-terminal residues | Reference |
| --- | --- | --- | --- | --- | --- |
| *Rhodothermus marinus* CdnE | 6E0K | I | 13 | 0 | Ref. 8 |
| *Elizabethkingia meningoseptica* CdnE | 6E0M | I | 6 | 1 | Ref. 8 |
| *Flavobacteriaceae* CdnE | 6WT8 | I | 13 | 4 | Ref. 5 |
| *Capnocytophaga granulosa* CdnE | 6WT9 | I | 17 | 0 | Ref. 5 |
| *V. cholerae* DncV | 4TXZ | II | 46 | 23 | Ref. 23 |
| *Enterobacter* CdnD | 7LJL | II | 66 | 27 | Ref. 24 |
| *Salmonella* CdnD | 7LJM | II | 66 | 27 | Ref. 24 |
| *Bacteroides fragilis* CdnB | 7LJO | II (short) | 11 | 50 | Ref. 24 |
| *Bradyrhizobium diazoefficiens* CdnG | 7LJN | II (short) | 7 | 49 | Ref. 24 |
| *E. coli* CdnC | 6P80 | III | 8 | 0 | Ref. 4 |
| *P. aeruginosa* CdnD | 6P82 | III | 5 | 0 | Ref. 4 |

**Supplementary Table S4.**
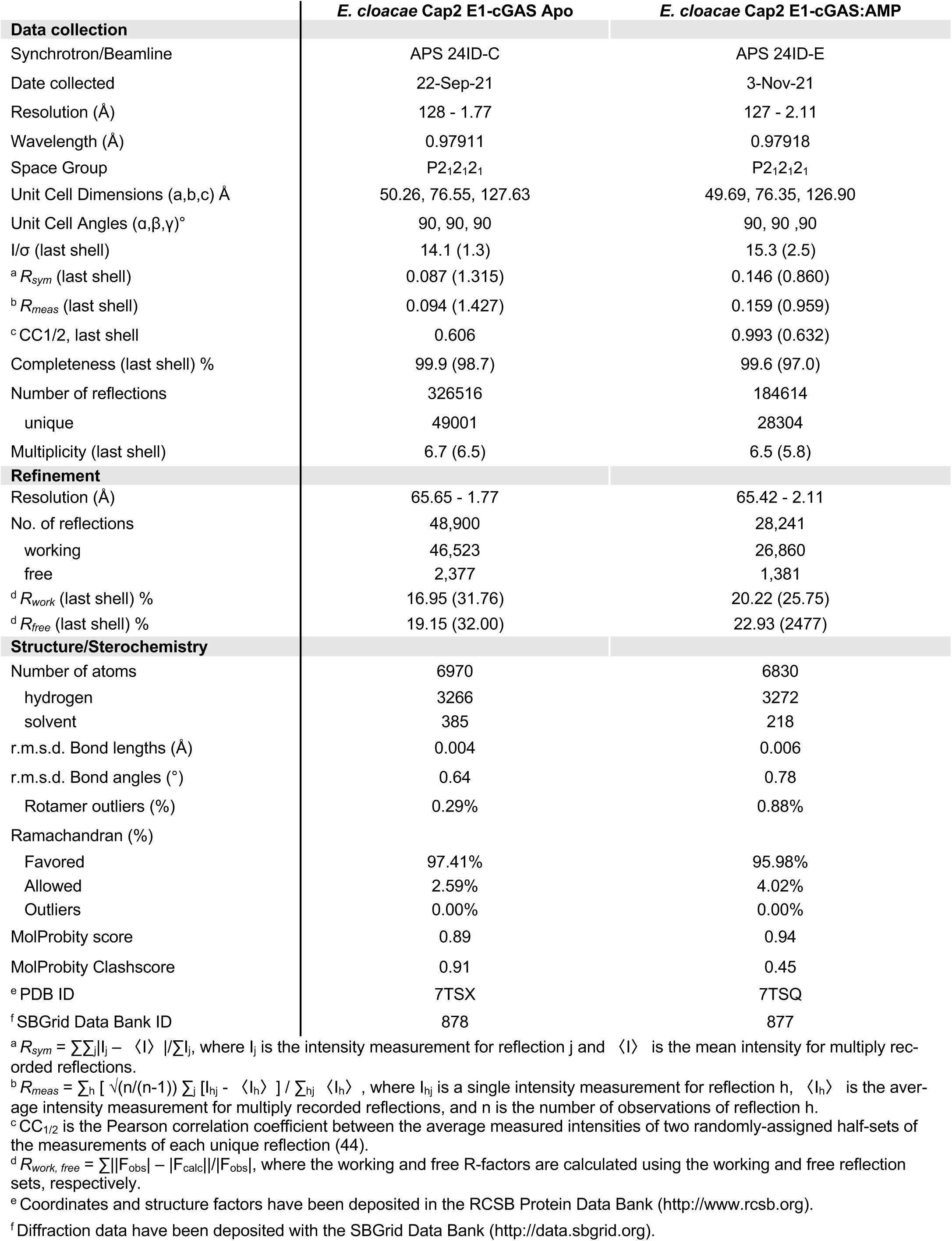
Crystallographic data collection and refinement.

|  | *E. cloacae* Cap2 E1-cGAS Apo | *E. cloacae* Cap2 E1-cGAS:AMP |
| --- | --- | --- |
| **Data collection** |  |  |
| Synchrotron/Beamline | APS 24ID-C | APS 24ID-E |
| Date collected | 22-Sep-21 | 3-Nov-21 |
| Resolution (Å) | 128 - 1.77 | 127 - 2.11 |
| Wavelength (Å) | 0.97911 | 0.97918 |
| Space Group | P2<sub>1</sub>2<sub>1</sub>2<sub>1</sub> | P2<sub>1</sub>2<sub>1</sub>2<sub>1</sub> |
| Unit Cell Dimensions (a,b,c) Å | 50.26, 76.55, 127.63 | 49.69, 76.35, 126.90 |
| Unit Cell Angles (α,β,γ)° | 90, 90, 90 | 90, 90, 90 |
| I/σ (last shell) | 14.1 (1.3) | 15.3 (2.5) |
| <sup>a</sup> *R*<sub>sym</sub> (last shell) | 0.087 (1.315) | 0.146 (0.860) |
| <sup>b</sup> *R*<sub>meas</sub> (last shell) | 0.094 (1.427) | 0.159 (0.959) |
| <sup>c</sup> CC1/2, last shell | 0.606 | 0.993 (0.632) |
| Completeness (last shell) % | 99.9 (98.7) | 99.6 (97.0) |
| Number of reflections | 326516 | 184614 |
| unique | 49001 | 28304 |
| Multiplicity (last shell) | 6.7 (6.5) | 6.5 (5.8) |
| **Refinement** |  |  |
| Resolution (Å) | 65.65 - 1.77 | 65.42 - 2.11 |
| No. of reflections | 48,900 | 28,241 |
| working | 46,523 | 26,860 |
| free | 2,377 | 1,381 |
| <sup>d</sup> *R*<sub>work</sub> (last shell) % | 16.95 (31.76) | 20.22 (25.75) |
| <sup>d</sup> *R*<sub>free</sub> (last shell) % | 19.15 (32.00) | 22.93 (24.77) |
| **Structure/Stereochemistry** |  |  |
| Number of atoms | 6970 | 6830 |
| hydrogen | 3266 | 3272 |
| solvent | 385 | 218 |
| r.m.s.d. Bond lengths (Å) | 0.004 | 0.006 |
| r.m.s.d. Bond angles (°) | 0.64 | 0.78 |
| Rotamer outliers (%) | 0.29% | 0.88% |
| Ramachandran (%) |  |  |
| Favored | 97.41% | 95.98% |
| Allowed | 2.59% | 4.02% |
| Outliers | 0.00% | 0.00% |
| MolProbity score | 0.89 | 0.94 |
| MolProbity Clashscore | 0.91 | 0.45 |
| <sup>e</sup> PDB ID | 7TSX | 7TSQ |
| <sup>f</sup> SBGrid Data Bank ID | 878 | 877 |

<sup>a</sup> $R_{sym} = \sum_j |I_j - \langle I \rangle| / \sum_j I_j$ , where $I_j$ is the intensity measurement for reflection $j$ and $\langle I \rangle$ is the mean intensity for multiply recorded reflections.
<sup>b</sup> $R_{meas} = \sum_h [ \sqrt{(n/(n-1))} \sum_j |I_{hj} - \langle I_h \rangle| ] / \sum_h \langle I_h \rangle$ , where $I_{hj}$ is a single intensity measurement for reflection $h$ , $\langle I_h \rangle$ is the average intensity measurement for multiply recorded reflections, and $n$ is the number of observations of reflection $h$ .
<sup>c</sup> CC<sub>1/2</sub> is the Pearson correlation coefficient between the average measured intensities of two randomly-assigned half-sets of the measurements of each unique reflection (44).
<sup>d</sup> $R_{work, free} = \sum [ |F_{obs}| - |F_{calc}| ] / |F_{obs}|$ , where the working and free R-factors are calculated using the working and free reflection sets, respectively.
<sup>e</sup> Coordinates and structure factors have been deposited in the RCSB Protein Data Bank (http://www.rcsb.org).
<sup>f</sup> Diffraction data have been deposited with the SBGrid Data Bank (http://data.sbgrid.org).

**Supplementary Table S5.** Trypsin Mass Spectrometry of Cap3-treated cGAS-GFP.

| Peptide | Peak Area | Relative Peak Area |
| --- | --- | --- |
| **Untreated *E. cloacae* cGAS-GFP (peptides ending in VSK.G)** |  |  |
| K.ISSTMVSGGIGSGSNGSSGSVSK.G | 8.45E+07 |  |
| K.ISSTMVSGGIGSGSN(+.98)GSSGSVSK.G | 3.27E+09 |  |
| K.ISSTM(+15.99)VSGGIGSGSNGSSGSVSK.G | 7.86E+06 |  |
| **Cap3-treated *E. cloacae* cGAS-GFP (peptides ending in VSK.G)** |  |  |
| R.FAGIGSGSN(+.98)GSSGSVSK.G | 2.58E+08 | 0.495 |
| **A.GIGSGSN(+.98)GSSGSVSK.G** | **5.21E+08** | **1.000** |
| G.IGSGSN(+.98)GSSGSVSK.G | 8.89E+05 | 0.002 |
| I.GSGSN(+.98)GSSGSVSK.G | 7.16E+05 | 0.001 |
| I.GSGSNGSSGSVSK.G | 1.48E+05 | 0.000 |
| **Untreated *V. cholerae* cGAS-GFP (peptides ending in VSK.G)** |  |  |
| K.ISSTMVSGGIGSGSNGSSGSVSK.G | 8.45E+07 |  |
| K.ISSTMVSGGIGSGSN(+.98)GSSGSVSK.G | 3.27E+09 |  |
| K.ISSTM(+15.99)VSGGIGSGSNGSSGSVSK.G | 7.86E+06 |  |
| **Cap3-treated *V. cholerae* cGAS-GFP (peptides ending in VSK.G)** |  |  |
| K.ISSTM(+15.99)VSGGIGSGSN(+.98)GSSGSVSK.G | 1.03E+08 | 0.054 |
| S.STM(+15.99)VSGGIGSGSN(+.98)GSSGSVSK.G | 6.25E+07 | 0.033 |
| S.TM(+15.99)VSGGIGSGSN(+.98)GSSGSVSK.G | 4.78E+06 | 0.003 |
| T.M(+15.99)VSGGIGSGSN(+.98)GSSGSVSK.G | 5.85E+08 | 0.306 |
| M.VSGGIGSGSN(+.98)GSSGSVSK.G | 5.37E+07 | 0.028 |
| S.GGIGSGSN(+.98)GSSGSVSK.G | 1.88E+07 | 0.010 |
| **G.GIGSGSN(+.98)GSSGSVSK.G** | **1.91E+09** | **1.000** |
| G.IGSGSN(+.98)GSSGSVSK.G | 9.15E+05 | 0.000 |
| I.GSGSN(+.98)GSSGSVSK.G | 2.95E+06 | 0.002 |

**Supplementary Table S6.** *E. coli* strains and plasmids used in this study.

| Strain # | *E. coli* | Plasmid # | Plasmid details |
| --- | --- | --- | --- |
| AW2413 | Omnipir | N/A | N/A; strain used for plasmid construction/cloning |
| AW2482 | Omnipir | pAW1382 | pLOCO2-*gfp* |
| AW2668 | Omnipir | pAW1575 | pLOCO2-*dncV* operon v2 (VSP-I) *V. cholerae* C6706, *Vc* operon 1 |
| AW2779 | Omnipir | pAW1601 | pLOCO2-*dncV* operon, *capV*_C62S |
| AW2783 | Omnipir | pAW1593 | pLOCO2-*dncV* operon, *dncV*_DID141AIA |
| AW2864 | Omnipir | pHL0108 | pLOCO2-*dncV* operon, $\Delta cap2$ |
| AW2865 | Omnipir | pHL0004 | pLOCO2-*dncV* operon, $\Delta cap3$ |
| AW2866 | Omnipir | pHL0033 | pLOCO2-*dncV* operon, *cap2*-3xFLAG $\Delta cap3$ |
| AW2867 | Omnipir | pHL0103 | pLOCO2-*dncV* operon, *cap2*_R390A-3xFLAG $\Delta cap3$ |
| AW2868 | Omnipir | pHL0085 | pLOCO2-*dncV* operon, *cap2*-3xFLAG_C522A $\Delta cap3$ |
| AW2869 | Omnipir | pHL0084 | pLOCO2-*dncV* operon, *cap2*-3xFLAG_C90A $\Delta cap3$ |
| AW2870 | Omnipir | pHL0104 | pLOCO2-*dncV* operon, *dncV*_G436E *cap2*-3xFLAG $\Delta cap3$ |
| AW2871 | Omnipir | pHL0105 | pLOCO2-*dncV* operon, *dncV*_SG435-6EE *cap2*-3xFLAG $\Delta cap3$ |
| AW2872 | Omnipir | pHL0114 | pLOCO2-*dncV* operon, *capV*_C62S *dncV*-int1vsvG *cap2*-3xFLAG $\Delta cap3$ |
| AW2873 | Omnipir | pHL0117 | pLOCO2-*dncV* operon, *capV*_C62S *dncV*_DID141AIA-intvsvG1 *cap2*-3xFLAG $\Delta cap3$ |
| AW2874 | Omnipir | pHL0116 | pLOCO2-*dncV* operon, *capV*_C62S *dncV*-int1vsvG *cap2*_C522A-3xFLAG $\Delta cap3$ |
| AW2875 | Omnipir | pHL0043 | pLOCO2-*dncV* operon, *dncV*-intvsvG-1 |
| AW2784 | Omnipir | pAW1608 | pTACxc-*sfGFP* |
| AW2787 | Omnipir | pAW1611 | pTACxc-*cap3* |
| AW2876 | Omnipir | pHL0061 | pTACxc-*cap3*_H101A H103A |
| AW2017 | MG1655 | N/A | N/A; strain used for phage and IP/ELISA experiments |
| N/A | Rosetta2 pLysS | pKDC4233 | *V. cholerae* His6-Cap3 |
| N/A | Rosetta2 pLysS | pKDC4246 | *V. cholerae* His6-DncV (for anti-cGAS antibody generation) |
| N/A | LOBSTR | pKDC4268 | *V. cholerae* His6-DncV-GFP |
| N/A | LOBSTR | pKDC4293 | *V. cholerae* His6-DncV(M433E)-GFP |
| N/A | LOBSTR | pKDC4294 | *V. cholerae* His6-DncV(M433A)-GFP |
| N/A | LOBSTR | pKDC4295 | *V. cholerae* His6-DncV(V434E)-GFP |
| N/A | LOBSTR | pKDC4296 | *V. cholerae* His6-DncV(M433A/V434A)-GFP |
| N/A | LOBSTR | pKDC4297 | *V. cholerae* His6-DncV(S435E)-GFP |
| N/A | LOBSTR | pKDC4298 | *V. cholerae* His6-DncV(G436E)-GFP |
| N/A | Rosetta2 pLysS | pKDC4334 | *V. cholerae* His6-Cap3 E38A |
| N/A | Rosetta2 pLysS | pKDC4336 | *V. cholerae* His6-Cap3 D113A |
| N/A | LOBSTR | pKDC4430 | *E. cloacae* His6-CdnD02-GFP |
| N/A | LOBSTR | pKDC4454 | *E. cloacae* His6-CdnD02(Q375A)-GFP |
| N/A | LOBSTR | pKDC4455 | *E. cloacae* His6-CdnD02(K376A)-GFP |
| N/A | LOBSTR | pKDC4456 | *E. cloacae* His6-CdnD02(T377A)-GFP |
| N/A | LOBSTR | pKDC4457 | *E. cloacae* His6-CdnD02(G378E)-GFP |
| N/A | LOBSTR | pKDC4458 | *E. cloacae* His6-CdnD02(R379A)-GFP |
| N/A | LOBSTR | pKDC4459 | *E. cloacae* His6-CdnD02(F380A)-GFP |
| N/A | LOBSTR | pKDC4460 | *E. cloacae* His6-CdnD02(A381E)-GFP |
| N/A | Rosetta2 pLysS | pKDC4483 | *E. cloacae* His6-Cap2/CdnD02 |
| N/A | Rosetta2 pLysS | pKDC4521 | *E. cloacae* His6-Cap3 |
| N/A | Rosetta2 pLysS | pKDC4534 | *E. cloacae* His6-Cap2(C548A)/CdnD02 |
| N/A | Rosetta2 pLysS | pKDC4539 | *E. cloacae* His6-Cap2/CdnD02(T377A) |
| N/A | Rosetta2 pLysS | pKDC4540 | *E. cloacae* His6-Cap2/CdnD02(G378E) |
| N/A | Rosetta2 pLysS | pKDC4541 | *E. cloacae* His6-Cap2/CdnD02(R379A) |
| N/A | Rosetta2 pLysS | pKDC4542 | *E. cloacae* His6-Cap2/CdnD02(F380A) |
| N/A | Rosetta2 pLysS | pKDC4543 | *E. cloacae* His6-Cap2/CdnD02(A381E) |
| N/A | Rosetta2 pLysS | pKDC4546 | *E. cloacae* His6-Cap2(C109A)/CdnD02 |
| N/A | Rosetta2 pLysS | pKDC4547 | *E. cloacae* His6-Cap2(C109A/C548A)/CdnD02 |
| N/A | Rosetta2 pLysS | pKDC4550 | *E. cloacae* His6-Cap2(R420A)/CdnD02 |
| N/A | Rosetta2 pLysS | pKDC4551 | *E. cloacae* His6-Cap2(D479A)/CdnD02 |
| N/A | Rosetta2 pLysS | pKDC4552 | *E. cloacae* His6-Cap2(D411A/N417A)/CdnD02 |
| N/A | Rosetta2 pLysS | pKDC4580 | *E. cloacae* HA-CdnD02 |
| N/A | Rosetta2 pLysS | pKDC4582 | *E. cloacae* HA-CdnD02/Cap2 |
| N/A | Rosetta2 pLysS | pKDC4584 | *E. cloacae* His6-Cap2 |
| N/A | Rosetta2 pLysS | pKDC4593 | *E. cloacae* His6-Cap2/CdnD02 (1-362) |
| N/A | Rosetta2 pLysS | pKDC4604 | *E. cloacae* HA-CdnD02/Cap2 C109A/C548A |
| N/A | Rosetta2 pLysS | pKDC4629 | *E. cloacae* His6-Cap3 E54A |
| N/A | Rosetta2 pLysS | pKDC4630 | *E. cloacae* His6-Cap3 D128A |
| N/A | Rosetta2 pLysS | pKDC4659 | *E. cloacae* His6-CdnD02/Cap2 (C109A/C548A) (for cryoEM) |
| N/A | Rosetta2 pLysS | pKDC4809 | *E. cloacae* His6-Cap2(363-600,C548A)-GSG-CdnD02(370-381) (for crystallography) |
| N/A | Rosetta2 pLysS | pKDC4811 | *E. cloacae* His6-Cap2(374-600,C548A)-GSG-CdnD02(370-381) (for crystallography) |
| N/A | Rosetta2 pLysS | pKDC4841 | *E. cloacae* His6-Cap2(R369A)/CdnD02 |
| N/A | Rosetta2 pLysS | pKDC4843 | *E. cloacae* His6-Cap2(I546A/W549A)/CdnD02 |
| N/A | Rosetta2 pLysS | pKDC4852 | *E. cloacae* His6-CdnD02/Cap2(C109A/C548A/T517K) |
| N/A | Rosetta2 pLysS | pKDC4854 | *E. cloacae* His6-CdnD02(R29A/R34A)/Cap2(C109A/C548A) |
| N/A | Rosetta2 pLysS | pKDC4855 | *E. cloacae* His6-CdnD02(T30K)/Cap2(C109A/C548A) |
| N/A | Rosetta2 pLysS | pKDC4856 | *E. cloacae* His6-CdnD02(W170A/L171A)/Cap2(C109A/C548A) |

**Supplementary Table S7.** Protein sequences.

| Organism | Gene ID | Accession Number |
| --- | --- | --- |
| *Vibrio cholerae* C6706 | *capV* | WP_001133548.1 |
| *Vibrio cholerae* C6706 | *dncV* (cGAS) | WP_001901330.1 |
| *Vibrio cholerae* C6706 | *cap2* | WP_001884104.1 |
| *Vibrio cholerae* C6706 | *cap3* | WP_001286069.1 |
| *Enterobacter hormaechei* subsp. *hoffmannii* UCI 50 | *cap4* | WP_032676399.1 |
| *Enterobacter hormaechei* subsp. *hoffmannii* UCI 50 | *cdnD02* (cGAS) | WP_032676400.1 |
| *Enterobacter hormaechei* subsp. *hoffmannii* UCI 50 | *cap2* | WP_050010101.1 |
| *Enterobacter hormaechei* subsp. *hoffmannii* UCI 50 | *cap3* | WP_157903655.1 |

**Supplementary Table S8.** Phages used in this study.

| Phage Name | Propagation strain | Reference and source |
| --- | --- | --- |
| T2 | MG1655 | CGSC12141 |
| T4 | MG1655 | CGSC12143 |
| T5 | MG1655 | CGSC12144 |
| T6 | MG1655 | NCCB 3461 |

